# Patient-Specific Midbrain Organoids with CRISPR Correction Recapitulate Neuronopathic Gaucher Disease Phenotypes and Enable Evaluation of Novel Therapies

**DOI:** 10.1101/2025.11.06.686937

**Authors:** Yi Lin, Benjamin Liou, Venette Fannin, Stuart Adler, Christopher N. Mayhew, Jason E. Hammonds, Yueh-Chiang Hu, Jason Tchieu, Wujuan Zhang, Xueheng Zhao, Rebecca L. Beres, Kenneth DR. Setchell, Ahmet Kaynak, Xiaoyang Qi, Ricardo A. Feldman, Ying Sun

## Abstract

Neuronopathic Gaucher disease (nGD) is a lysosomal storage disorder caused by *GBA1* mutations, leading to defective acid β-glucosidase (GCase) and accumulation of glycosphingolipid substrates, causing inflammation and neurodegeneration. Patients with nGD manifest severe neurological symptoms, but current animal models fail to fully recapitulate human condition, posing a major barrier to the development of effective therapies targeting the brain. To bridge this gap, we have developed midbrain-like organoids (MLOs) from human induced pluripotent stem cells (hiPSCs) of nGD patients with *GBA1*^L444P/P415R^ and *GBA1*^L444P/RecNcil^ mutations to model nGD brain pathogenesis. These nGD MLOs exhibited GCase deficiency, resulting in diminished enzymatic function, accumulation of lipid substrates, widespread transcriptomic changes, and impaired dopaminergic neuron differentiation, mirroring nGD pathology. *GBA1* mutation correction mediated by CRISPR/Cas9 restored GCase activity, normalized lipid substrate levels, and rescued dopaminergic neuron function, confirming the causal role of *GBA1* mutations during early brain development. Using this novel platform, we further evaluated therapeutic strategies, including SapC-DOPS nanovesicles delivering GCase, AAV9-GBA1 gene therapy, and substrate reduction therapy with GZ452, a glucosylceramide synthase inhibitor currently under clinical investigation. These treatments either restored GCase activity, reduced lipid substrate accumulation, improved autophagic and lysosomal abnormalities, or ameliorated dysregulated genes involved in neural development. These patient-specific, 3D neural models offer a transformative, physiologically relevant platform for unravelling disease mechanisms and accelerating the discovery of therapies for patients with nGD.

## Introduction

Gaucher disease (GD) is a lysosomal storage disorder caused by *GBA1* mutations, which impair acid β-glucosidase (GCase). Defective GCase leads to accumulation of its glycosphingolipid substrates, glucosylceramide (GluCer) and glucosylsphingosine (GluSph), triggering inflammation and neurodegeneration. GD affects 1/450 in Ashkenazi individuals and 1/50,000 globally ^1^. There are three GD types: Type 1 present mainly visceral symptoms and is not life-threatening ^1^. while neuronopathic GD (nGD, Types 2 and 3) involves severe neurological manifestations. Type 2 patients typically die by age 2, indicating early central nervous system (CNS) defects ^2,3^.

Current knowledge of brain pathogenesis in nGD is limited to a few postmortem brain analyses showing prominent neuronal loss and astrogliosis in the midbrain ^4–6^. GD mouse models, including GCase knockout, irreversible GCase inhibitor conduritol B-epoxide (CBE) induced mouse models, double transgenic mutant model harboring both *Gba1* and Saposin C mutations (Saposin C is the activator of GCase), have been widely used to study disease pathogenesis and therapies ^7–12^. These models replicate neuronal loss and astrogliosis ^4–6,8,13^, but have limitations. The knockout and CBE-induced models lack *GBA1* mutations, and the double transgenic model has a second gene that may change the GD phenotype, and knock-in models fail to mimic human phenotypes ^10,14^. For example, mice with the *Gba1*^L444P/L444P^ or *Gba1*^D409H/D409H^ mutations show no neuronopathic disease despite <10% of residual GCase activity ^10,14–16^. These constraints highlight the need for physiologically relevant human nGD models.

Brain organoids derived from human induced pluripotent stem cells (hiPSC) emerged as a powerful 3D model to study brain development and CNS diseases ^17^. Unlike 2D monolayer neural cells, brain organoids contain diverse neural and glial cell types arranged in 3D comprising subventricular zone and multiple organized cellular layers that more closely represents the spatial architecture of the human brain ^18^. They have been used to study neurodegenerative and lysosomal diseases, including Parkinson’s disease (PD), Alzheimer’s disease, and Sandhoff disease ^19–23^. A brain organoid derived from hiPSCs of a healthy individual with *GBA1* knockout and α-synuclein overexpression exhibited characteristic PD markers ^24^. However, this model lacks patient-specific *GBA1* mutations needed for clinical relevance. To develop a relevant model, we developed novel midbrain organoids from nGD patients with *GBA1* mutations, a model not yet well studied.

The midbrain region is prominently affected in nGD ^4–6^. Patients with nGD often exhibit impaired vertical gaze and movement disorders. These symptoms correlate with midbrain involvement due to the sensitivity of this region to neuroinflammatory and degenerative processes ^4,25^. Both human and mouse studies indicate that the midbrain shows prominent substrate accumulation compared with other brain regions, suggesting it is particularly susceptible to pathological burden in GD midbrain ^4,26–28^. To model the midbrain-specific disease, we derived midbrain-like organoids (MLOs) from Type 2 patient-derived hiPSCs carrying either *GBA1*^L444P/P415R^ or *GBA1*^L444P/RecNcil^ mutations. These GD MLOs showed reduced GCase activity, lipid substrate accumulation, transcriptomic alterations, and impaired dopaminergic neuron differentiation. CRISPR/Cas9-mediated correction of *GBA1* mutation confirmed the causal role of *GBA1* mutations in these phenotypes and mitigated disease phenotypes. To explore therapeutic potential, we tested three emerging treatments in GD MLOs: CNS accessible enzyme therapy (SapC-DOPS-GCase), gene therapy (AAV9-GBA1) and substrate reduction therapy (GZ452). Our results demonstrate the utility of patient-derived MLOs for studying nGD pathogenesis and advance drug development.

## Materials and Methods

### Maintenance of hiPSCs and generation of MLOs

Three hiPSCs lines (WT-75.1, GD2-1260 and GD2-10-257) generated previously ^29,30^ (Supplementary Table 1) were maintained in mTeSR1 complete medium (StemCell) on six-well plates coated with Vitronectin (1:100 diluted in DPBS, 1 mL/well, incubated at RT for 1 hour). HiPSCs at ∼70% confluency were treated and replated at 1:20 to 1:60 ratios, with optional 10 μM ROCK inhibitor (Y-27632, StemCell) for survival. All hiPSC lines were authenticated using short tandem repeat (STR) profiling and were verified as mycoplasma-free.

MLO generation method was modified based on published protocol ^31,32^. Briefly, hiPSCs were pretreated with 50 μM Y-27632 for 1 hour, dissociated with Accutase (Millipore-Sigma) into singlets, centrifuged (200 × g, 5 minutes), and resuspended in embryoid body (EB) medium (EBM; consisting of DMEM/F12, 20% KnockOut Serum Replacement (KSR), 1% penicillin/streptomycin, GlutaMAX, NEAA, 55 μM β-mercaptoethanol, 1 μg/mL heparin, 3% FBS, 4 ng/mL bFGF, 1× CEPT). For each organoid, 1.2∼1.5 × 10⁴ hiPSC cells were seeded in 100 μL EBM per well in ultra-low attachment (ULA) U-bottom 96-well plates, centrifuged at 500 × g for 3 minutes, and incubated at 37°C, 5% CO₂; on day 2, 150 μL EBM with 4 ng/mL bFGF (no ROCK inhibitor) was added. Once the EBs grow to 400 to 475 μm in size, they were transferred to a new ULA 96-well plate with 125 μL brain organoid generation medium (BGM, consisting of 50% DMEM/F12, 50% Neurobasal, 1× N2, 1× B27 without vitamin A, 1% penicillin/streptomycin, 1% GlutaMAX, 1% NEAA, 55 μM β-mercaptoethanol, 1 μg/mL heparin) supplemented with dual SMAD inhibitors (SMADi: 2 μM dorsomorphin, 2 μM A83-01, 3 μM CHIR99021, 1 μM IWP2), refreshed every other day until day 11. On day 7 or 8, mesencephalic floor plate induction was initiated by adding 125 μL BGM containing 1× SMADi, 100 ng/mL FGF8, and 2 μM SAG until day 16. On day 11, four EBs were transferred to one well of ultra-low attachment 24-well plates with 500 μL BGM-MLOs induction medium, consisting of BGM with 100 ng/mL FGF8, 2 μM SAG, 200 ng/mL laminin, 2.5 μg/mL insulin, 2% Growth Factor Reduced (GFR) Matrigel, and refreshed with Matrigel on day 14. On day 16, three MLOs per well were transferred to ultra-low attachment 6-well plates in brain organoid maturation medium (BGM-BMM), consisting of BGM with 10 ng/mL BDNF, 10 ng/mL GDNF, 200 μM ascorbic acid, 125 μM db-cAMP, 1% GFR Matrigel, cultured on an orbital shaker (100 rpm) to reduce spontaneous fusion with medium changes every 3 days. Starting at day 30, GFR Matrigel was removed, and brain organoids were cultured solely in BGM medium till the date of analysis.

### Immunostaining and image analysis of sectioned MLOs

Organoids were washed with 1xDPBS and fixed in 4% paraformaldehyde (PFA) overnight at 4°C. After washing with PBS-T (1 x DPBS with 0.1% Tween 20), organoids were cryoprotected in 30% sucrose solution until fully equilibrated. Samples were then embedded in gelatin solution (7.5% gelatin, 10% sucrose in 1xDPBS) at 37°C for 1 hour, snap-frozen in a dry ice/ethanol slurry, and stored at -80°C. Cryosectioning was performed to obtain 16-μm-thick sections. For immunofluorescence, sections were air-dried and outlined with a hydrophobic PAP pen before blocked with 5% normal goat serum in PBS-T for 1 hour, incubated overnight with primary antibodies (Supplementary Table 1) in primary antibody dilution buffer (PBS-T with 5% BSA and 0.05% sodium azide), and followed by secondary antibodies (Alexa Fluor® conjugates) for 1 hour at room temperature. Slides were washed, stained with DAPI solution (0.2 μg/mL), mounted with Antifade Mounting Medium (VECTASHIELD®, H-1000-10), and imaged at 10× magnification for whole organoid scans or 60× magnification for detailed cellular analysis using a fluorescence confocal microscope (Nikon Eclipse *Ti*) (Nikon, Tokyo, Japan) to assess neuronal organization of MLOs. The morphologies of neurons, astrocytes, and colocalization analysis were performed by taking multi-layer stacking (Z-stack) images under 60× oil immersion objective lens. Quantitative immunofluorescence analyses (e.g., cell counts for FOXP1+, FOXG1+, SOX2+ and Ki67+ cells, as well as marker colocalization) were performed using ImageJ (NIH) on at least 3 - 5 randomly selected non-overlapping fields of view (FOVs) per organoid section, with a minimum of 3 organoids per differentiation batch. Each FOV was imaged at consistent magnification (60x) and z-stack depth to ensure comparable sampling across conditions. Data from individual FOVs were first averaged within each organoid to obtain an organoid-level mean, and then biological replicates (independent differentiations, n ≥ 3) were averaged to generate the final group mean ± SEM.

### Genome editing patient iPSC clones

The iPSC line (GD2-1260) used for gene correction was derived from fibroblasts obtained from a GD Type 2 patient carrying compound heterozygous *GBA1* mutations (P415R/L444P) ^29^. A guide (g) RNA was designed to target SpCas9-mediated double strand break introduction proximal to the L444P mutation (*GBA1* mutation at nt14446, T>C) in GD2-1260. Oligonucleotides (caccGAAGAACGACCCGGACGCAG and aaacCTGCGTCCGGGTCGTTCTTC; overhangs in lower case; target mutation site underlined; IDT) transcribing the sgRNA target sequence were annealed and cloned via BbsI restriction digest into plasmid pX458M that contains a U6 promoter-driven sgRNA and a SpCas9-2A-EGFP expression cassette. The pX458M plasmid is modified from the pX458 plasmid (addgene #48138) ^33^ and carries an optimized sgRNA scaffold ^34^. The targeting activity of this plasmid was validated by T7E1 assay using 293 cells. GD2-1260 cells were transfected with pX458M and a phosphorothioate-modified single stranded oligonucleotide (ssODN; IDT) donor template using TransIT-LT1 (Mirus). The ssODN (Supplementary Figure 1A) was designed to introduce the desired wildtype *GBA1* sequence flanked by homology arms to the targeted genomic region. The ssODN was also designed to contain silent mutations to prevent retargeting by SpCas9 and to introduce a BtgI restriction site to facilitate identification of targeted clones. Forty-eight hours post-transfection, GFP-positive cells were isolated by FACS and replated at cloning density (250-500 cells/well of a 6 well matrigel-coated plate). Replated cells were cultured for 4 days in mTeSR1 containing 10% CloneR (StemCell Technologies), using the manufacturer’s recommended protocol. Cells were subsequently fed daily with mTeSR1 for an additional 9 days before colonies were manually harvested and expanded for genotyping. Primers VS4247 (gtgcgtaactttgtcgacagtcc) and VS4249 (ctgagagtgtgatcctgccaag) were used to PCR amplify the targeted *GBA1* genomic region and products were subjected to BtgI digestion to identify putative edited clones. Selected clones were subsequently confirmed by Sanger sequencing. This gene editing strategy is expected to also target the *GBA1* pseudogene due to the identical target sequence which limits the gene correction on certain mutations (e.g., P415R) ^35,36^, but the chance to target other off-targets is low due to low off-target scores ranked based on the MIT Specificity Score analysis ^37^.

### SapC-DOPS-fGCase treatment of MLOs

As previously described ^38,39^, SapC-DOPS nanovesicles were formulated by combining saposin C (SapC) with dioleoylphosphatidylserine (DOPS) to encapsulate either recombinant human acid β-glucosidase (fGCase, Freeline Therapeutics) or fluorescent CellVue Maroon (CVM) for uptake studies. The SapC-DOPS-fGCase complex was prepared at a final concentration of 0.6 µg/mL fGCase and added to the BGM culture medium of MLOs derived from WT-75.1, GD2-1260, and GD2-10-257 hiPSC lines. To assess nanovesicle uptake, WT-75.1 MLOs at week 13 were co-cultured with SapC-DOPS-CVM for 48 hours, followed by confocal imaging to confirm internalization of the fluorescent CVM within the organoids.

For enzyme delivery and therapeutic evaluation, MLOs were treated with SapC-DOPS-fGCase or SapC-DOPS alone (control) for a 48-hour period to assess uptake and enzyme activity. For long-term treatment efficacy, MLOs were treated for two weeks before evaluating therapeutic efficacy. During treatment, MLOs were maintained in BGM media containing nanovesicles and were replaced every 3 days to ensure consistent exposure. Post-treatment, MLOs were harvested for confocal imaging, enzymatic assays, and biochemical analyses to assess GCase activity, protein expression, and correction of GD phenotypes.

### AAV injection into MLOs

Transfer vector of AAV9-GBA1 virus containing GBA1/GFP expressing cassette (CB-GBA1-IRES-GFP) was packaged in AAV9 capsid at AAVnerGene (Rockville, MD, USA). For the delivery of AAV9-GBA1 gene therapy to MLOs, a precise injection protocol was employed using a nanoliter injector system (World Precision Instruments). MLOs derived from WT-75.1, GD2-1260, and GD2-10-257 hiPSC lines were placed in a dish containing sterile 1 x PBS. A glass pipette connected to a gas injector was lowered into the center of each MLO at a desired depth of 200 to 300 µm, as determined by microscopic visualization. A volume of 500 nL (0.5µL) of AAV9-GBA1 vector, at a concentration of 3.6 × 10¹³ vg/mL, was injected into ten sites of MLOs at a volume of 50 nL per injection site. The injection glass needle was retracted slowly after completion of the injection to minimize damage and ensure vector distribution. Morphology and viability were assessed post-injection, and MLOs were then maintained in culture for 3 weeks prior to subsequent analyses.

### Substrate Reduction Therapy treatment of MLOs

For evaluation of the tolerated dose of Substrate Reduction Therapy (SRT), WT-75.1 MLOs were treated with SRT compound GZ452 (AstaTech, P14969), also named GZ-682452, an analogue of venglustat ^40^, at concentrations of 0.3, 1, 2, and 3 µM starting on day 0 for a period of 6 weeks. MLO size was measured weekly using bright-field microscopy and analyzed with ImageJ software to assess morphological changes. For short-term SRT treatment, 13-week-old GD2-1260 MLO were treated with 300nM GZ452 for 3 weeks, with medium refreshed every 3 days to maintain consistent drug exposure. For long-term SRT treatment, GZ452 treatment at the concentration of 300nM was started at the beginning of MLO generation (day 1) and continued for 28 weeks before collection for analysis. Untreated MLOs served as controls.

### Glycosphingolipids analyses

MLOs were washed with ice-cold PBS and homogenized and sonicated in sterile ddH_2_O. Glycosphingolipids were then extracted from tissue homogenates using chloroform/methanol following protocol outlined previously ^41^. Aliquots of lipids extracts were processed for GluCer and GluSph quantification by ultra-high-performance chromatography coupled to tandem mass spectrometry (UHPLC-MS/MS) using a Waters Xevo TQ-S Micro triple quadrupole mass spectrometer (Waters, Milford, MA) at CCHMC Clinical Mass Spectrometry Laboratory. Chromatographic separation for GluCer and GluSph was achieved using a XSelect CSH C18 XP Column (100 x 2.1 mm, 2.5 µm, Waters) column. Quantification by LC-MS/MS was operated in the multiple reaction monitoring (MRM) mode, with detection of the transition pair of the individual protonated parent ions of GluCers and daughter ion m/z 264.2. GluSph was measured by monitoring the mass transition m/z 462.3> 282.4. The sphingoid base for all GluCer species analyzed is d18:1. Calibration curves were prepared for C16 GluCer, C18 GluCer, C24 GluCer, and C24:1 GluCer using C18 glucosyl(ß) ceramide-D5 as the internal standard. Quantification of GluCer species with various fatty acid chain lengths was realized by the calibration curve of each species or with the closest fatty acyl chain length. The quantification of GluSph was based on the calibration curve using glucosyl(β) sphingosine-D5 as internal standard. The calibration curve for GluCer and GluSph was 25 pg–10 ng on column. Three QCs at low, medium and high levels (50 pg, 0.5 ng and 5.0 ng) were prepared in organic solvent and analyzed along with samples. The GluCer and GluSph levels in MLO were normalized to total MLO protein (mg) that were used for glycosphingolipids analyses. Protein mass was determined by BCA assay and glycosphingolipid was expressed as pmol/mg protein. Additionally, GluSph levels in the culture medium were quantified and normalized to the medium volume (pmol/mL) ^42^.

### Measuring Dopamine levels by ELISA

Dopamine levels in MLO culture medium were quantified using the Dopamine ELISA Kit (Abnova) that involves dopamine extraction, acylation and enzymatic conversion before assay. Briefly, MLO culture medium was collected from 4 MLOs cultured in 3 mL BGM medium for 72 hours. Dopamine extraction was performed by pipetting 10 µL of standards, controls, and 750 µL of the sample into the extraction plate wells, filling each to 750 µL with deionized water, followed by 25 µL of TE buffer. Shake the covered plate for 60 minutes at room temperature (RT, 20-25°C) at 600 rpm, wash with 1 mL wash buffer twice, then acylate with 150 µL acylation buffer and 25 µL acylation reagent for 20 minutes. After washing again, elute with 100 µL hydrochloric acid, and transfer 90 µL of supernatant to the microtiter plate for enzymatic conversion with 25 µL freshly prepared enzyme solution, incubating at 37°C for 2 hours. Then, 100 µL of the converted samples and 50 µL dopamine antiserum were added to the dopamine microtiter strips. The mixture was then incubated overnight at 2-8°C. The strips were then washed and incubated with 100 µL enzyme conjugate for 30 minutes, followed by 100 µL of substrate for 20-30 minutes, and finally, 100 µL of stop solution was added. Absorbance was measured at 450 nm within 10 minutes, and dopamine concentrations were calculated using a calibration curve, applying the correction factor (10/500 = 0.02) to adjust and normalize by sample volume.

### GCase activity assay

GCase enzyme activities in hiPSCs and MLOs were determined fluorometrically with 4-methylumberlliferyl-β-D-glucopyranoside (4MU-Glc) in the presence of the GCase irreversible inhibitor, Conduritol B epoxide (CBE, 2 mM, Millipore, Bedford, MD) as previously described ^43^. Briefly, hiPSC cell pellet or MLO tissues were homogenized using Precellys Evo Homogenizer/CK Mix beads (Bertin Technologies, France) in 1% sodium taurocholate/1%Triton X-100 (Tc/Tx) solution. GCase enzyme activity was then determined by 4MU-Glc as substrate in 0.25% Tc/Tx diluted in 0.1 M citrate phosphate buffer (pH 5.6). Protein concentrations were determined by BCA assay for normalization. The GCase-specific activity was calculated by subtracting non-specific activity (with CBE) from total activity (without CBE) and normalized to total protein mass.

### Transcriptome analysis of MLOs

Total RNAs were extracted from week 8 MLOs using the RNeasy Mini Kit (Qiagen). Total RNAs (150 to 300 ng), quantified by Qubit (Invitrogen) high-sensitivity spectrofluorometric assay, were poly-A selected and reverse transcribed using Illumina’s TruSeq® stranded mRNA library preparation kit (Illumina). Each sample was fitted with one of 96 adapters containing different 8-base molecular barcodes for high-level multiplexing. After 15 cycles of PCR amplification, completed libraries were sequenced on NovaSeq 6000 (Illumina), generating 30 million high-quality 100-base-long paired-end reads per sample. A quality control check on the fastq files was performed using FastQC. Upon passing basic quality metrics, the reads were trimmed to remove adapters and low-quality reads using default parameters in Trimmomatic (Version 0.33). The trimmed reads were then mapped to human reference genome GRCh38 using default parameters with strandness option in Hisat2 (Version 2.0.5), achieving a mapping rate greater than 90%. Transcript or gene abundance was determined using kallisto (Version 0.43.1). A transcriptome index in kallisto was created using Ensembl cDNA sequences for the reference genome. This index was then used to quantify transcript abundance in raw counts and transcript per million (TPM). Differential expression analysis was performed using the R package DESeq2 ^44^. Genes with BaseMean ≥ 50, |fold change| ≥ 2, p-adj ≤ 0.05 were identified as differentially expressed genes (DEGs). Heatmaps were made using the R package ggplot2. Gene enrichment analysis using cellular component, KEGG pathways, molecular function, and biological process (GO analysis) was performed using DAVID 6.8 ^45^. Three MLOs were pooled for each sample and three samples were profiled for each genotype. MLO Sequencing datasets and partially processed results have been deposited to the National Center for Biotechnology Information (NCBI)’s Gene Expression Omnibus (GEO) database (GSE303993).

### Gene expression analysis by qRT-PCR

Total RNA was isolated from MLO tissue using the RNeasy Kit (Qiagen). Quantitative RT-PCR was carried out as described previously using primers listed in Supplementary Table 1 ^46^. Housekeeping gene *GAPDH* or *ACTB* was used as internal controls for mRNA quantification. Relative expression of mRNAs was determined by the 2^-ΔΔ^CT method.

### Immunoblotting

Immunoblotting was used to detect protein expression in iPSCs and MLOs. Briefly, brain organoids or cells were homogenized in freshly prepared RIPA buffer with protease inhibitors and anti-phosphatase inhibitors using a Precellys Evo Homogenizer (Bertin). The samples were then centrifuged at 12,000 g for 10 min to produce lysate for electrophoresis, membrane transferring, and detection using primary and secondary antibodies indicated in Supplementary Table 1. Semi-quantification of proteins was performed using ImageJ (NIH).

### Statistical analysis

For comparisons between two groups, data were analyzed using unpaired two-tailed Student’s t-tests when the sample size was ≥ 6 per group and normality was confirmed by the Shapiro-Wilk test. When the normality assumption was not met or when sample sizes were small (n < 6), the non-parametric Mann-Whitney U test was used instead. For comparisons involving three or more groups, one-way ANOVA followed by Tukey’s multiple comparison test was applied when data were normally distributed; otherwise, the non-parametric Dunn’s multiple comparison test was used. Exclusion of outliers was made based on cutoffs of the mean ±2 standard deviations. All statistical analyses were performed using GraphPad Prism 10 software. Exact p-values are reported throughout the manuscript and figures where feasible. A p-value < 0.05 was considered statistically significant.

## Results

### Generation and characterization of MLOs derived from hiPSCs

To generate MLOs from iPSCs, healthy hiPSCs were differentiated following a stepwise protocol, neural induction, neuroepithelial expansion, and maturation, adopted with modifications from previous studies ^31,32^ (Figure 1A). Midbrain specification during development requires appropriate posterior differentiation signals ^47,48^. Our protocol used sequential activation/inhibition of critical signaling pathways involved in midbrain specification: Wnt signaling (activated by CHIR99021), BMP signaling (inhibited by dual-SMAD), and FGF signaling (activated by FGF8) (Figure 1A). MLOs were matured and characterized using immunofluorescence, gene expression, and immunoblotting. Week 8 MLOs showed diverse neural cell types, including pan-neurons (Tuj1/NeuN), astrocytes (GFAP), dopaminergic (DA) neurons (FOXA2/TH), and neural progenitor cells (SOX2/Ki67) (Figure 1B). Comparative analysis of forebrain marker FOXG1 expression between MLOs and cerebral organoids (COs) showed barely detectable FOXG1 in MLOs (Figure 1C). Quantitative RT-PCR analysis demonstrated a significant increase in midbrain/DA neuron-specific genes (*FOXA2*, *ASCL1*, *LMX1A*, *PLZF*, and *TH*) from week 3 and week 8 MLOs, demonstrating the midbrain identity of these organoids. Expression of glial-specific genes (*GLAST*, *S100B*) increased over time, while pluripotency markers (*SOX2*, *NANOG*, *OCT4*) were downregulated, suggesting progressive neural differentiation (Figure 1D). Immunoblot further validated hiPSCs differentiation into MLOs by expression of neuronal (Tuj1, MAP2), DA neuronal (TH), and glial (GFAP, S100B) markers at week 8, with reduced Sox2 expression in MLOs versus hiPSCs (Figure 1E). Together, these results confirm the successful generation of MLOs with midbrain-like cellular composition and molecular identities.

**Figure 1.**
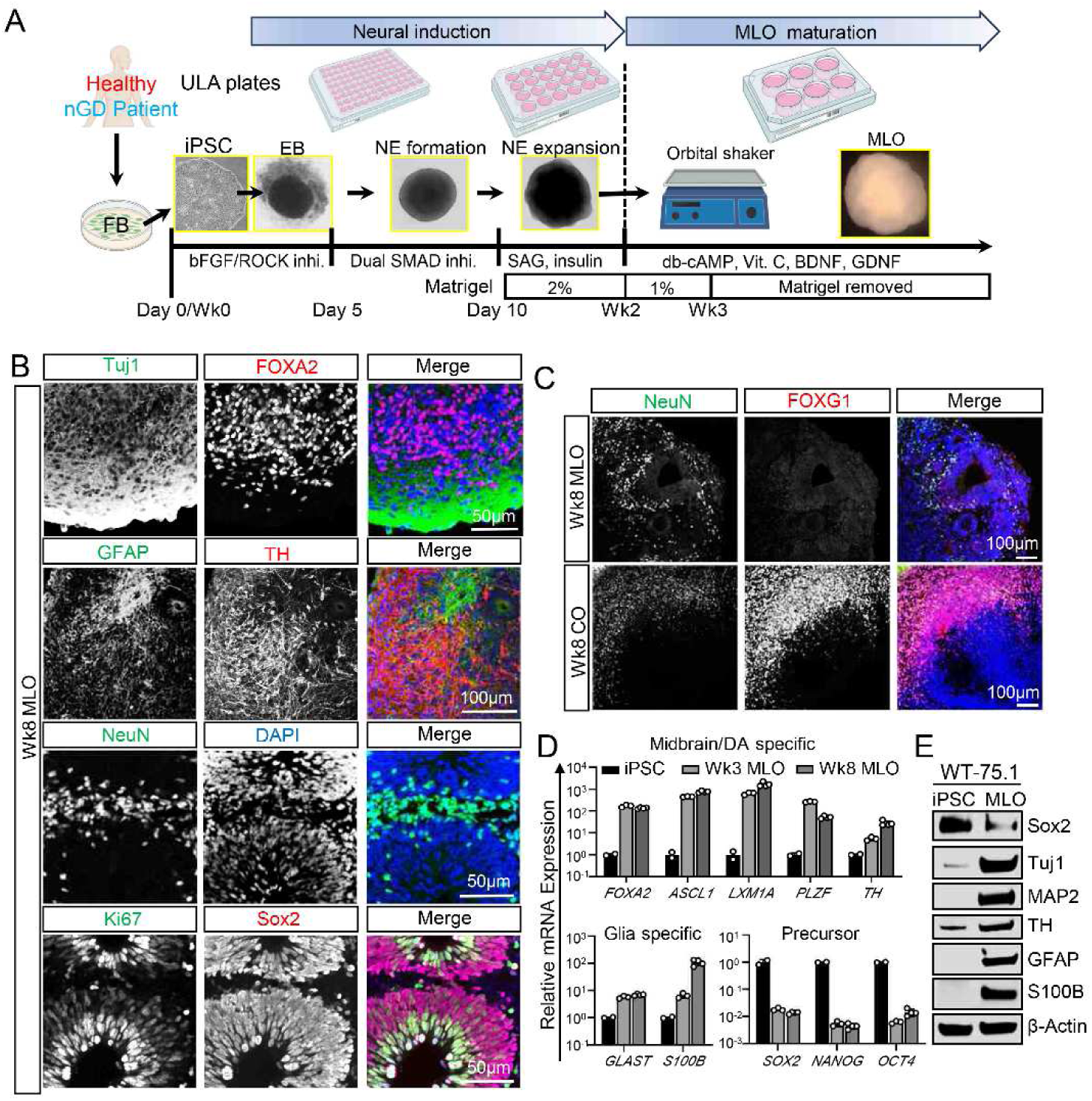
Generation and characterization of midbrain-like organoids (MLOs) from healthy human (h) iPSCs (WT-75.1 hiPSCs). (A) A schematic overview of the procedures for generating MLOs from healthy WT-75.1 hiPSCs. (B) Representative confocal images showing architectural structure of week (Wk) 8 MLOs containing new-born neurons (Tuj1/NeuN), astrocytes (GFAP), dopaminergic neurons (FOXA2/TH) and neural progenitor cells (SOX2/Ki67). Merge images show the distribution of those cell markers. (C) FOXG1 expression in MLO and cerebral organoid (CO). Transcription factor FOXG1 (forebrain marker) was enriched in CO at Wk8 of differentiation but absent in MLO. Pan-neurons (NeuN) were both present in MLO and CO, as shown by NeuN immunostaining (neuronal marker). (D) Quantitative analysis of cell type specific genes expression for midbrain/dopaminergic neuron (*FOXA2*/*ASCL1*/*LXM1A*/*PLZF*/*TH*), glial cells (*GLAST*/*S100B*) and multipotent stem cells (*SOX2*/*NANOG*/*OCT4*) in Wk3 and Wk8 MLOs (n=3 MLOs pooled for each group) by qRT-PCR. (E) Immunoblot of Sox2, Tuj1, MAP2, TH, GFAP and S100B in WT-75.1 hiPSCs and its derived MLO (Wk8, n=3 MLOs pooled for each group) lysate. β-Actin was used as a loading control.

### GCase deficiency in GD MLOs leads to glycosphingolipids accumulation and altered transcriptomic profiling

To model nGD and investigate the impact of GCase deficiency on MLOs, we generated GD MLO using the GD2-1260 hiPSC line derived from GD Type 2 patient harboring *GBA1*^L444P/P415R^, compound heterozygous mutations, using same protocol in Figure 1 (Figure 2) ^29^. Midbrain identity of GD MLOs was confirmed by TH/FOXA2 expression (Figure 3). Immunoblot showed an approximately 85% reduction in GCase protein levels in week 8 GD2-1260 MLOs versus WT-75.1 MLOs (Figure 2A). Consistent with reduced GCase protein, GCase activity decreased to 14.2% in GD2-1260 hiPSCs and 15.0% in GD2-1260 MLOs vs. WT-75.1 (*p* < 0.001; Figure 2B). Bright-field imaging at weeks 4, 8, and 15 showed no difference on overall MLOs appearance between WT-75.1 and GD2-1260 MLOs (Figure 2C). MLO size measurements revealed a trend toward smaller GD2-1260 MLOs versus WT-75.1 at week 8, though not statistically significant (Figure 2D).

**Figure 2.**
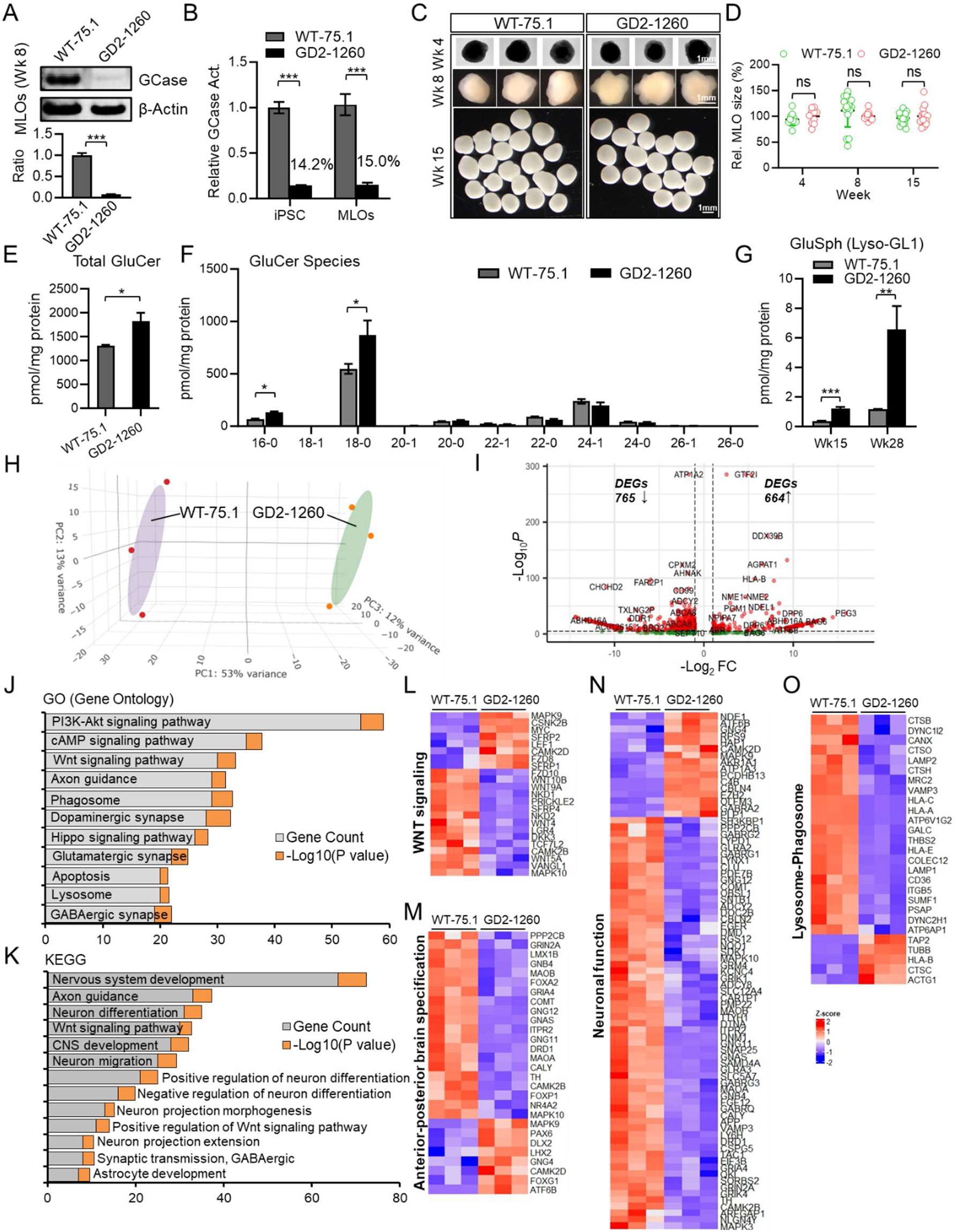
GCase deficiency drives glycosphingolipids accumulation and transcriptomic alteration in GD MLOs. The GD MLOs were generated as in Figure 1A. (A) Reduced GCase protein in GD MLO (GD2-1260). Wk 8 MLOs (n=3) were pooled as a biological sample. β-Actin was used as loading control. (B) GCase activity in hiPSCs and MLOs (> 3 MLOs were pooled for each group). Data was normalized to WT-75.1 control. (C) Representative images of WT-75.1 and GD2-1260 MLOs at Wks 4, 8 and 15 of differentiation. (D) MLO size was measured based on area of MLO spheres and normalized to WT-75.1 control at each indicated time points. N ≥ 10 MLOs were quantified per group. (E, F) Measurement of total glucosylceramide (GluCer) and GluCer species in Wk 15 MLO. (B, D, E. G) *p < 0.05, ***p < 0.001, ns, not significant, unpaired Student’s t test. (G) Glucosylsphingosine (GluSph) levels in Wk15 and Wk28 MLOs (3∼5 MLOs were pooled for each group). GluCer and GluSph levels in the organoids were measured by LC-MS/MS and normalized by corresponding total protein of MLO tissue lysate. (H) 3D Principal Component Analysis (PCA) of bulk RNA sequencing (RNA-seq) data. The Euclidean distance of the normalized gene expression among healthy control (WT-75.1) and GD (GD2-1260) MLOs was used for sample clustering. Ellipsoids around each group indicate the distribution and spread of the samples within the sample group. Wk 8 MLOs (n=3) were pooled as one biological sample, and three samples were profiled in each group. (I) MA plot showing the distinct genes differentially expressed in GD MLOs. Statistically significant differentially expressed genes (DEGs; |fold change|≥ 1, p-adj ≤ 0.05 and base mean ≥ 50) were highlighted in red. Number of DEGs downregulated and upregulated in GD2-1260 MLO against WT-75.1 MLO were shown. FC, fold-change. (J, K) Dysregulated pathway in GD MLOs analyzed by GO (gene ontology) (J) and Kyoto Encyclopedia of Genes and Genomes (KEGG) (K) enrichment of DEGs. Both gene counts and level of significance (-log10 of *p* value) were shown as stacked columns for each category. (L-O) Heat maps of dysregulated pathways or biological functions in GD MLO. Specifically, aberrant expressions of genes involved in WNT signaling (L), anterior-posterior brain specification (M), neuronal function (N) and lysosome-phagosome (O) were shown.

**Figure 3.**
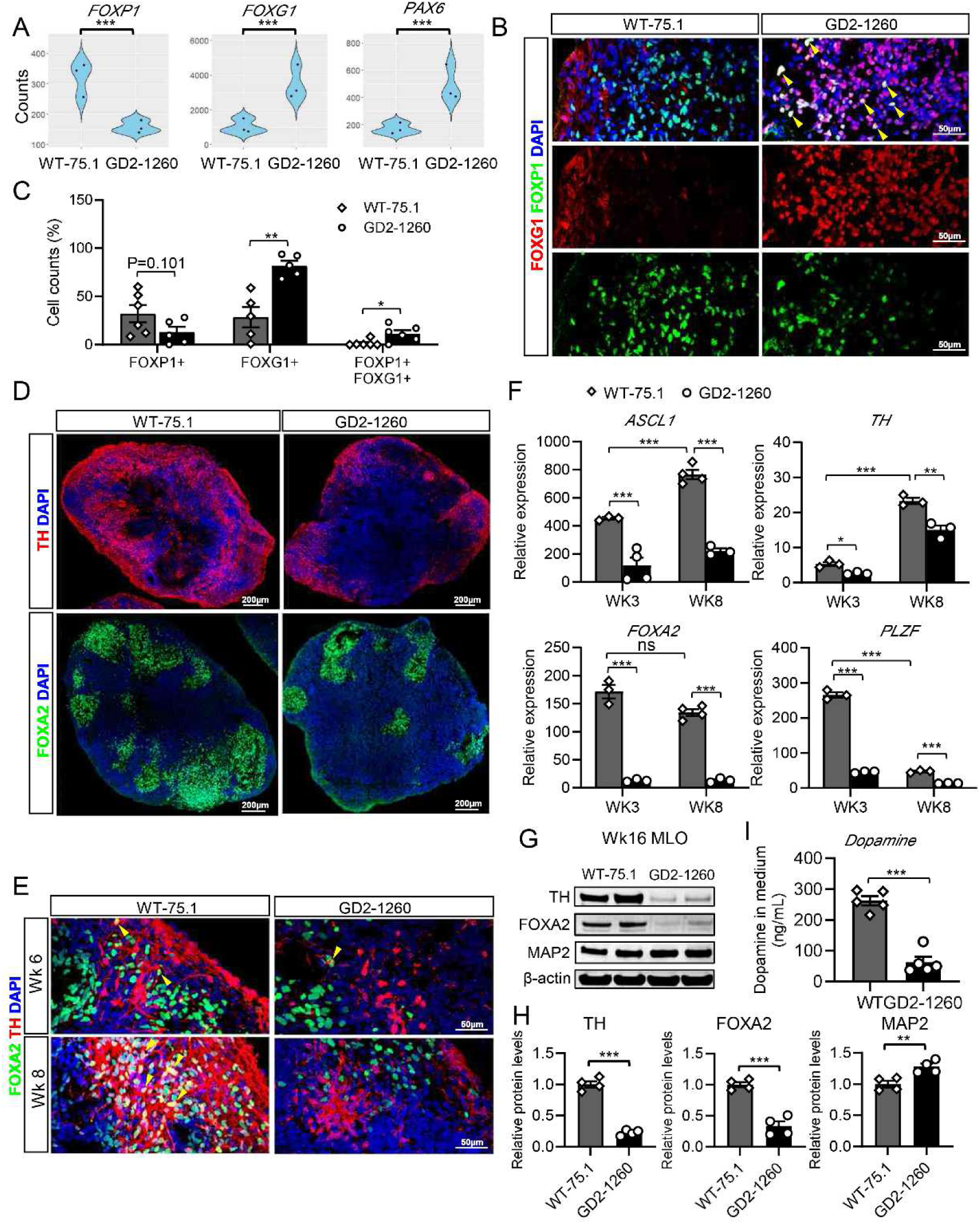
Skewed specification of midbrain patterning and dopaminergic neuron differentiation in GD MLOs. (A) Gene expression of *FOXP1*, *FOXG1*, and *PAX6* in week 8 WT-75.1 and GD2-1260 MLOs. Data were plotted using RNA sequencing counts. ***p < 0.001, unpaired Student’s t test. (B, C) Aberrant expression of FOXP1/FOXG1 transcription machinery for forebrain/midbrain patterning in GD MLOs. Representative confocal images (B) and quantification of Wk8 WT-75.1 and GD2-1260 MLOs, immunostained for FOXP1 (red) and FOXG1 (green), with DAPI (blue) labeling nuclei. Yellow arrows indicate FOXP1+FOXG1+ cells. Scale bar, 50 µm. *p < 0.05, **p < 0.01, unpaired Student’s t test. (D) Confocal images of Wk6 MLOs, immunostained for midbrain patterning markers FOXA2 (green) or TH (red), with DAPI (blue) labeling nuclei. Scale bar, 100 µm. (E) Representative images of differentiating DA neurons in MLOs derived from WT-75.1 and GD2-1260 hiPSCs. TH (red), FOXA2 (green) were co-stained, with DAPI (blue) labeling nuclei. Yellow arrows indicate TH+FOXA2+ cells. Scale bar, 50 µm. (F) Quantification of midbrain progenitor markers *ASCL1*, *TH*, *LMX1A*, and *PLZF* expression in WT-75.1 and GD2-1260 MLOs at Wk3 and Wk8, measured by qRT-PCR and normalized to WT-75.1 hiPSC cells. Data are presented as mean ± SEM (n = 3-4 MLOs per group). *p < 0.05, **p < 0.01. (G) Immunoblot analysis of midbrain/dopaminergic neuron markers TH, FOXA2, and MAP2 in Wk16 MLOs. Protein samples were extracted from n=3 MLOs from each group. β-Actin was used as a loading control. (H) Relative protein levels of TH, FOXA2, and MAP2 in Wk8 GD2-1260 MLOs compared to WT-75.1. *p < 0.05, **p < 0.01, unpaired Student’s t test. (I) Dopamine levels in MLO culture medium assay by ELISA. Culture medium from 4 GD2-1260 MLOs or WT-75.1 MLOs at Wk12 cultured in 3 mL BGM medium for 72 hours was assayed. Data are presented as mean ± SEM (n = 5 per group). ***p < 0.001, unpaired Student’s t test.

Loss of GCase function in GD disrupts glycosphingolipids metabolism, a primary pathogenic factor in nGD ^49^. Mass spectrometry analysis demonstrated significant glycosphingolipid substrate accumulation in GD2-1260 MLOs. At week 15, total glucosylceramide (GluCer) level was elevated approximately 1.39-fold in GD2-1260 MLOs compared to WT-75.1 MLOs (p < 0.05; Figure 2E). GluCer analysis revealed marked accumulation of species 18:0 and 16:0, particularly brain predominant species (18:0) in GD2-1260 MLOs, while other species remained largely unchanged (Figure 2F, Supplementary Figure 2). Similarly, glucosylsphingosine (GluSph), a toxic lipid associated with nGD ^4,40,50^, was significantly elevated approximately 3.3-fold and 5.6-fold in GD2-1260 MLOs at weeks 15 and 28, respectively, versus WT-75.1 controls (Figure 2G, Supplementary Figure 2). These findings indicate profound lipid substrates dysregulation in GCase-deficient GD MLOs, consistent with the biochemical hallmark of GD.

To explore the transcriptomic consequences of GCase deficiency, we performed bulk RNA sequencing on week 8 MLOs, when DA neurons maturation and GD phenotypes were evident. Principal component analysis (PCA) revealed distinct clustering of WT-75.1 and GD2-1260 MLOs, with the first principal component accounting for 53% of the variance, indicating significant transcriptomic differences between two genotype groups (Figure 2H). An MA plot identified 1,429 differentially expressed genes (DEGs) in GD2-1260 MLOs compared to WT-75.1, with 664 genes upregulated and 765 genes downregulated. GO analysis of these DEGs revealed significant enrichment in pathways including cAMP, PI3K-AKT, and WNT signaling that control the nervous system development, axon guidance, and neuron differentiation (Figure 2J). KEGG analysis similarly identified dysregulation in neural signaling pathways, synaptic transmission, and lysosomal function (Figure 2K). The heatmap of these DEGs elucidated specific gene expression changes within key pathways (Figure 2L-2O), including WNT signaling (Figure 2L) ^30^, anterior-posterior brain specification (Figure 2M), neuronal function (Figure 2N), and lysosome-phagosome (Figure 2O) in GD2-1260 MLOs, with many genes exhibiting upregulation or downregulation consistent with nGD pathology.

Collectively, these results demonstrate that *GBA1* mutation in GD MLOs leads to reduced enzyme activity, causes lipid substrate accumulation, and alters transcriptomic profiles, particularly in neural development, signaling, and lysosomal function. These findings provide insights into nGD mechanism and support MLOs as a model for studying nGD.

### Skewed specification of midbrain patterning and dopaminergic neuron differentiation in GD MLOs

Building on the molecular and transcriptomic hallmarks of GCase deficiency observed in nGD MLOs (Figure 2), we next investigated the impact on midbrain patterning and dopaminergic neuron differentiation (Figure 3). Firstly, RNA sequencing data revealed elevated FOXG1 and PAX6, and reduced FOXP1 mRNAs in GD2-1260 MLOs versus WT-75.1 MLOs at week 8 (Figure 2M, Figure 3A). These genes are fate-determining regulators in controlling proper anterior-posterior neural patterning ^51–53^. Immunofluorescence confirmed reduced FOXP1+ and increased FOXG1+ cell counts in GD2-1260 MLOs (Figure 3B, 3C). About 11.6% cells were FOXP1+ FOXG1+ cells in GD2-1260 MLOs, but they were barely seen in WT-75.1 MLOs. These results indicate a significant disruption in expression of genes involved in forebrain and midbrain patterning in GD MLOs.

The skewed specification of midbrain patterning may contribute to dysregulated dopaminergic neuron development in GCase-deficient GD MLOs (Figure 3). There were clear differences in the expression of midbrain-specific markers between WT-75.1 and GD2-1260 organoids at week 6 (Figure 3D). In WT-75.1 MLOs, FOXA2 (a midbrain progenitor marker) and TH (dopaminergic neuron marker) were strongly expressed and co-localized (Figure 3D, 3E), indicating normal midbrain patterning. In contrast, GD2-1260 MLOs showed markedly reduced FOXA2+, TH+ and FOXA2+TH+ cells (Figure 3D, 3E). Quantitative analysis confirmed the significant reduction of midbrain progenitor markers (*ASCL1*, *TH*, *FOXA2*, and *PLZF*) in GD2-1260 MLOs versus WT-75.1 (Figure 3F). At week 8, expression levels of *ASCL1*, *TH*, *FOXA2*, and *PLZF* were significantly reduced by approximately 71.0%, 44.4%, 90.0% and 69.8%, respectively, indicating a profound impairment in midbrain progenitor specification due to GCase deficiency. Immunoblot showed ∼77.5% and ∼66.5% reductions in TH and FOXA2, respectively, and a 27.7% increase in MAP2 in GD2-1260 versus WT-75.1 MLOs (Figure 3H). Dopaminergic function was determined by measuring dopamine levels in the culture medium (Figure 3I). Dopamine levels in week 12 MLOs culture medium (72 hours culture) were 76.0% (p < 0.001) lower in GD2-1260 MLOs versus WT-75.1 MLOs, confirming impaired dopaminergic function in GD MLOs.

These results indicate that GCase deficiency leads to skewed midbrain patterning and dopaminergic neuron differentiation, underscoring the critical role of GCase in midbrain development, as revealed for the first time, using patient iPSC-derived brain organoids.

### CRISPR/Cas9-mediated *GBA1* mutation correction rescues disease phenotypes in GD MLOs

GD is an autosomal recessive disorder. Individuals who are carriers or heterozygous for *GBA1* mutations typically do not exhibit disease phenotypes ^1^. To determine whether correction of the *GBA1* mutation could rescue GD phenotypes in GD MLOs, we used CRISPR/Cas9 technology to correct the L444P mutation in GD2-1260 hiPSCs (*GBA1*^L444P/P415R^) and generated isogenic iso-GD2-1260 hiPSCs with a heterozygous genotype of *GBA1*^WT/P415R^ (Figure 4A; Supplementary Figure 1). Karyotype analysis of GD2-1260 and iso-GD2-1260 demonstrated no detectable chromosomal abnormalities, confirming the genetic stability of both cell lines (Supplementary Figure 1D). Furthermore, genetic correction did not alter the expression of hiPSC related pluripotent genes (Supplementary Figure 1E). The MLOs derived from WT-75.1, GD2-1260, and iso-GD2-1260 hiPSCs were then compared to evaluating the impact of mutation correction on GD phenotypes.

**Figure 4.**
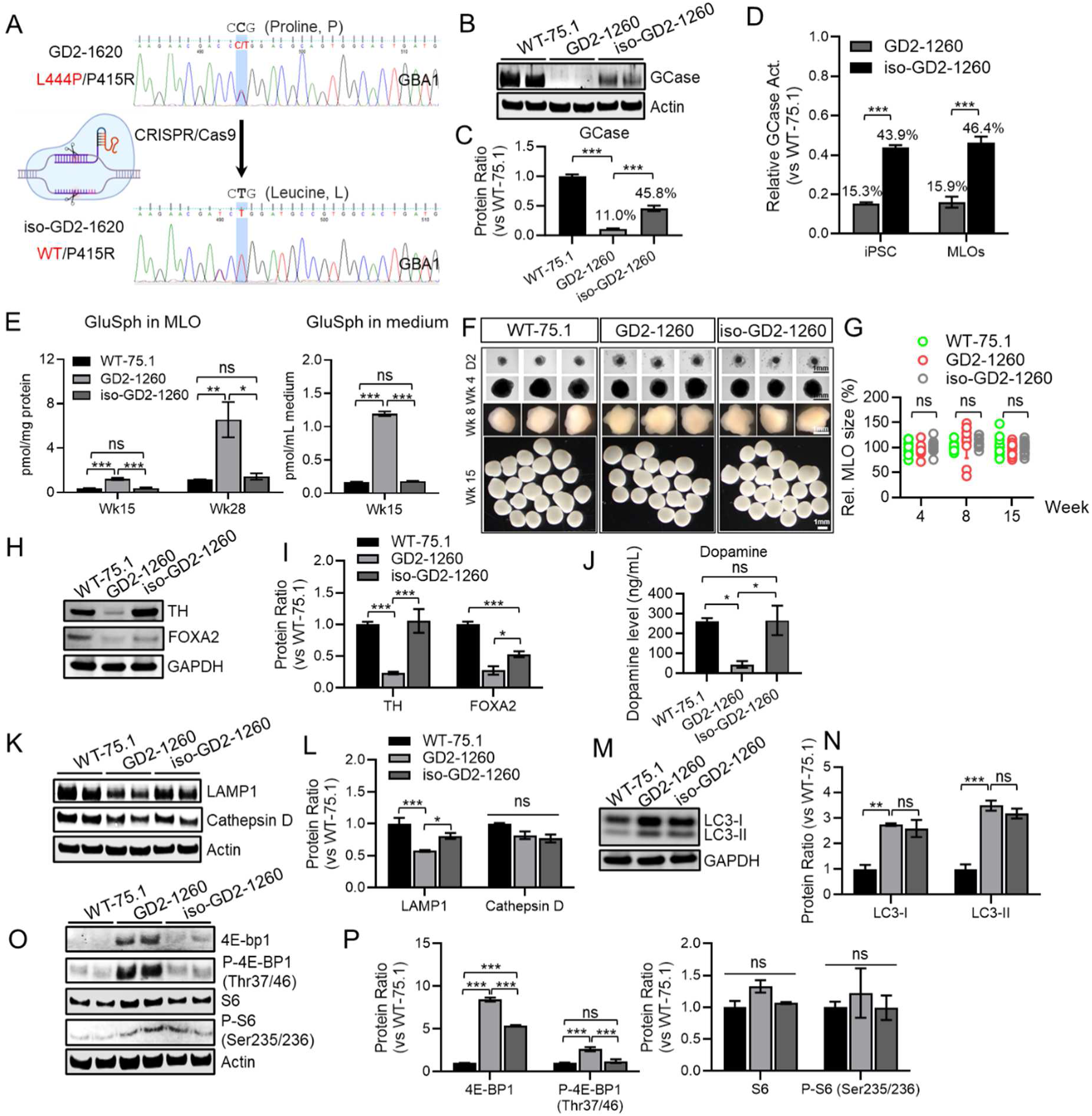
Mutation correction significantly rescued disease phenotypes in GD MLOs. (A) Schematic overview of CRISPR/Cas9-mediated mutation correction of the *GBA1* L444P mutation in GD2-1260 hiPSCs, converting the L444P (L444P/P415R) mutation (Proline, P to Leucine, L) to the wild-type sequence (WT-P415R), generating isogenic iso-GD2-1260 hiPSCs. The mutated base C in amino acid code ‘CCG’ for proline (P), was corrected to T to decode leucine (L, CTG), which was confirmed by genome sequencing of *GBA1* locus. (B, C) Immunoblot analysis of GCase protein and quantification in week 8 MLOs derived from WT-75.1, GD2-1260, and iso-GD2-1260 hiPSCs. β-Actin was used as a loading control. Data are presented as mean ± SEM (n = 2 pooled, and 3 biological replicates per group). ***p < 0.001. (D) Relative GCase activity in GD2-1260 and iso-GD2-1260 hiPSCs and Wk8 MLOs, normalized to WT-75.1 controls. Data are presented as mean ± SEM (2 MLOs pooled, n = 3 per group). ***p < 0.001. (E) Measurement of GluSph levels in WT-75.1, GD2-1260, and iso-GD2-1260 MLOs at Wk15 and Wk28 and their culture medium at Wk 15, quantified by LC-MS/MS and normalized to total protein of tissue lysate. Data are presented as mean ± SEM. For GluSph in MLO, three MLOs were pooled and n = 3 per group. For MLO secreted GluSph, MLO culture medium in wells containing four MLOs were collected, n = 3 per group. **p < 0.01; ns, not significant. (F) Representative bright-field images of WT-75.1, GD2-1260 and iso-GD2-1260 MLOs at Day 2, Wks 4, 8, and 15 of differentiation. Scale bar, 1 mm. For side-by-side comparison, images for WT-75.1 and GD2-1260 at Wks 4, 8 and 15 were taken from Fig 2C. (G) MLO size quantification for WT-75.1, GD2-1260 and iso-GD2-1260 MLOs at Wks 4, 8, and 15. MLOs size was analyzed by NIS elements and presented as the area (µm^2^) of MLO at indicated time point. N ≥ 10 MLOs were quantified per group. Data are presented as mean ± SEM. One-Way ANOVA, ns, not significant. (H, I) Immunoblot analysis of midbrain/dopaminergic neuron markers TH and FOXA2 (H) and their relative quantification (I) in Wk8 MLOs. Protein samples were extracted from n=3 MLOs from each group. GAPDH was used as a loading control. (J) Dopamine levels in the culture medium of Wk12 MLOs derived from WT-75.1, GD2-1260 and iso-GD2-1260 hiPSCs, measured after 72 hours in BGM medium (n = 4 MLOs per samples, 3 biological replicates). Data are presented as mean ± SEM (n = 5 per group). *p < 0.05, unpaired Student’s t test. (K, L) Immunoblot analysis of autophagy-lysosomal pathway markers LAMP1 and Cathepsin D (K) and quantification (L) in Wk16 MLOs. GAPDH was used as a loading control. Data are presented as mean ± SEM. (M, N) Immunoblot analysis of LC3-I and LC3-II (M) and quantification (N) in Wk16 MLOs. Protein samples were extracted from n=3 MLOs for each group. GAPDH was used as a loading control. (O) Immunoblot analysis of mTOR signaling pathway components [4E-BP1, P-4E-BP1(THR37/46), S6, and P-S6 (Ser235/236)] in Wk16 MLOs. β-Actin was used as a loading control. (P) Quantification of protein levels of mTOR signaling pathway components. Data are normalized to WT-75.1 and presented as mean ± SEM. Immunoblot analysis for panel H and I and K-P was performed using the lysate from 3 MLOs pooled per group, 3 repeated experiments. ***p < 0.001; ns, not significant. One-way ANOVA test with Tukey’s test.

At week 8, GCase protein in iso-GD2-1260 MLOs was restored to 45.8% of WT-75.1 levels (Figure 4B, 4C), consistent with correction of one *GBA1* allele ^29^. GCase activity also improved, reaching 43.9% and 46.4% of WT-75.1 levels in GD2-1260 hiPSCs and MLOs, respectively, versus approximate 15% in GD2-1260 hiPSCs and MLOs (Figure 4D). Despite reduced GCase, early neural rosette formation in GD2-1260 MLOs remained largely unaffected in GD2-1260 and iso-GD2-1260, with similar counts of SOX2+Ki67+ proliferating neural progenitor cells in neural rosettes (Supplementary Figure 3A, 3B), suggesting early neural development is not impacted by *GBA1* L444P mutation.

*GBA1* L444P mutation correction ameliorated lipid substrate accumulation in iso-GD2-1260 MLOs. GluSph levels which were elevated in GD2-1260 MLOs at weeks 15 and 28 (approximately 3.3-fold and 5.6-fold higher than WT-75.1, respectively; *p* < 0.01) was normalized to WT-75.1 levels (Figure 4E). Organoid size of iso-GD2-1260 MLOs was similar to WT-75.1 and GD2-1260 MLOs at week 4, 8 and 15 (Figure 4F, 4G).

Correction of the L444P mutation restored midbrain and dopaminergic neuron differentiation. TH and FOXA2 levels, which were reduced in week 16 GD2-1260 MLOs by approximately 76.4% and 72.4%, respectively, were restored in iso-GD2-1260 MLOs to WT-75.1 levels for TH, and approximately 53% of WT-75.1 levels for FOXA2 (Figure 4H, 4I). Dopamine levels in iso-GD2-1260 MLOs culture matched those in WT-75.1, unlike the reduced levels in GD2-1260 MLOs (Figure 4J), indicating restored dopaminergic function. The autophagy-lysosomal pathway, which is dysregulated in nGD, was partially corrected by *GBA1* mutation restoration (Figure 4K, 4L). Immunoblot analysis of lysosomal proteins showed that LAMP1 levels, which were significantly decreased in GD2-1260 MLOs by approximately 42.5% versus WT-75.1, were partially restored in iso-GD2-1260 MLOs (Figure 4K, 4L), however, Cathepsin D levels remained unchanged (Figure 4K, 4L). Dysregulated autophagy flux demonstrated by the elevated LC3-II and LC3-I was not significantly improved in isogenic MLOs compared to GD2-1260 MLOs (Figure 4M, 4N).

GCase deficiency leads to mTOR hyperactivation in nGD ^13,54,55^. In GD2-1260 MLOs, levels of 4E-BP1 and its phosphorylated form P-4E-BP1 (Thr37/46) were significantly elevated (Figure 4O). These levels were normalized or significantly reduced in iso-GD2-1260 MLOs. While S6 and its phosphorylated form (P-Ser235/236) remain unchanged (Figure 4O, 4P).

These results demonstrate that CRISPR/Cas9-mediated correction of *GBA1* mutation in GD2-1260 hiPSCs effectively rescues key nGD phenotypes and downstream effects, highlighting gene correction as a therapeutic strategy and validating MLOs as a preclinical disease model.

### SapC-DOPS nanoparticle-mediated GCase enzyme therapy corrects GD phenotypes in GD MLOs

SapC-DOPS nanoparticles, composed of saposin C (SapC) and dioleoylphosphatidylserine (DOPS), showed promise as a CNS-targeted delivery system for lysosomal disorders like nGD ^38,39^. The effectiveness of this approach in rescuing GD phenotypes was evaluated in MLOs derived from GD2-1260 hiPSCs and another GD2 hiPSC line, GD2-10-257, which carries the *GBA1*^L444P/RecNcil^ mutation ^56^. MLOs were derived from these hiPSCs following the protocol in Figure 1 and described in the Methods section. SapC-DOPS nanoparticle was formulated with a fGCase, a recombinant GCase variant with over 21-fold longer active half-life at lysosomal pH than wild-type GCase ^57,58^ (Figure 5A).

**Figure 5.**
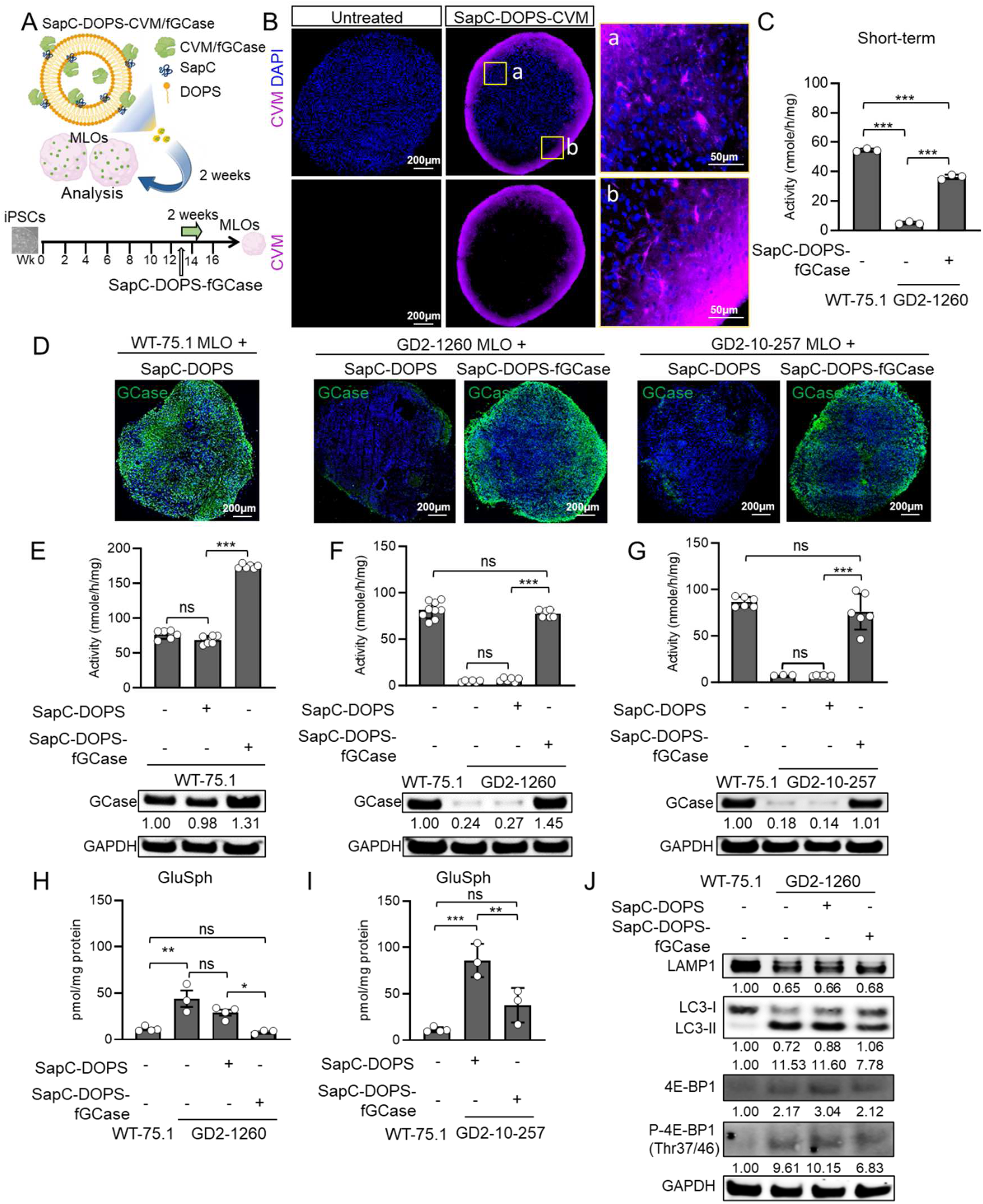
Delivery of GCase to MLOs via SapC-DOPS nanoparticles corrects GD phenotypes. (A) Schematic illustration of SapC-DOPS nanoparticle-mediated delivery of recombinant GCase (fGCase) to MLOs. SapC-DOPS nanoparticles carrying fGCase or fluorescent label CVM were cocultured with MLOs, followed by short-term (48 hours) or 2-week treatment period before analysis. (B) Confocal images of untreated and SapC-DOPS-CVM-treated MLOs, showing uptake of CVM (magenta) with DAPI (blue) labeling nuclei. Scale bars: 200 µm (left panel), 50 µm (right panels, magnified regions a and b). (C) GCase activity in WT-75.1 and GD2-1260 MLOs following a 48-hour treatment with SapC-DOPS-fGCase. Data are presented as mean ± SEM (3 MLOs pooled, n = 3 per group). ***p < 0.001. (D) Confocal images of WT-75.1, GD2-1260, and GD2-10-257 MLOs treated with SapC-DOPS or SapC-DOPS-fGCase for 2 weeks, immunostained for GCase (green) with DAPI (blue) labeling nuclei. Scale bar, 200 µm. (E-G) GCase activity and protein in WT-75.1 and GD (GD2-1260, GD2-10-257) MLOs treated with SapC-DOPS or SapC-DOPS-fGCase for 2 weeks, measured by enzymatic assay and immunoblot. Data are presented as mean ± SEM (3 MLOs pooled, n = 3∼4 per group). ***p < 0.001; ns, not significant. Protein samples were extracted from n=3 MLOs for each group. (H, I) GluSph levels in WT-75.1 and GD (GD2-1260, GD2-10-257) MLOs treated with SapC-DOPS or SapC-DOPS-fGCase for 2 weeks, quantified by LC-MS/MS and normalized to total protein. Data are presented as mean ± SEM (3 MLOs pooled, n = 3-4 per group). ***p < 0.001; **p < 0.01; ns, not significant. (J) Immunoblot analysis of autophagy-lysosomal and mTOR pathway proteins in SapC-DOPS or SapC-DOPS-fGCase treated GD2-1260 MLOs. GAPDH was used as a loading control. Protein samples were extracted from n=3 MLOs for each group. Protein levels are normalized to WT-75.1 untreated controls (set to 1.0).

To test SapC-DOPS uptake, WT-75.1 MLOs were co-cultured with CVM (CellVue Maroon) loaded SapC-DOPS nanoparticles for 48 hours. CVM signals were detected throughout the organoids, confirming successful cargo delivery (Figure 5B). SapC-DOPS-fGCase was then prepared with 0.6 µg/mL fGCase ^38^ and cocultured with MLOs. After 48 hours, treatment significantly increased GCase activity in GD2-1260 MLOs, restoring it to approximately 66.7% of WT-75.1 levels, confirming efficient delivery (Figure 5C).

To evaluate efficacy, GD2-1260 and GD2-10-257 MLOs were treated for two weeks with SapC-DOPS or SapC-DOPS-fGCase. Confocal imaging confirmed restored GCase expression, with SapC-DOPS-fGCase-treated GD2-1260 and GD2-10-257 MLOs showing increased GCase approaching the level in WT-75.1 MLO (Figure 5D). Additionally, SapC-DOPS-fGCase treatment significantly elevated GCase load in both dopaminergic neurons (TH+) and astrocytes (GFAP+), as shown by colocalization of GCase with TH and GFAP in GD2-1260 and GD2-10-257 MLOs, respectively (Supplementary Figure 4A-D). Consistent with the results obtained with GD2-1260 MLOs (Figure 4H), reduction of TH in GD2-10-257 MLOs at weeks 16 and 28 were evident (Supplementary Figure 4E). Corresponding to increased GCase protein, SapC-DOPS-fGCase treatment elevated GCase activity to 2.3-fold of untreated WT-75.1 MLO (Figure 5E). In GD2-1260 MLOs, SapC-DOPS-fGCase fully restored GCase activity to WT levels, while SapC-DOPS alone had no effect (Figure 5F). Similar GCase restoration was observed in the additional patient organoids, GD2-10-257 MLOs, indicating the effectiveness of this approach across different GD MLO models with various *GBA1* mutations (Figure 5G). Importantly, SapC-DOPS-fGCase significantly reduced elevated GluSph levels in GD2-1260 and GD2-10-257 MLOs, matching WT-75.1 levels (Figure 5H, 5I), confirming effective lipid substrate clearance by delivered fGCase.

Furthermore, we evaluated the impact of SapC-DOPS-fGCase on the autophagy and lysosomal pathway and mTOR signaling. fGCase was found effectively delivered to lysosomal compartments, evidenced by colocalization of LAMP1 and LC3-II in treated GD2-1260 and GD2-10-257 MLOs (Supplementary Figure 5A-D), indicating restoration of GCase in lysosomal and autophagosome compartments. However, analysis of protein levels showed that decreased LAMP1 expression in GD2-1260 MLOs was not altered following SapC-DOPS-fGCase treatment (Figure 5J). The elevated LC3-II levels, an indicator of impaired autophagic flux, were reduced upon treatment, suggesting enhanced autophagic activity (Figure 5J). Moreover, phosphorylated 4E-BP1 (Thr37/46), but not total 4E-BP1, was improved in SapC-DOPS-fGCase treated MLOs, reflecting a decrease in mTOR hyperactivation (Figure 5J). We anticipate that a longer duration of SapC-DOPS-fGCase exposure in nGD MLOs may produce a more robust therapeutic effect in rescuing nGD-associated phenotypes, which will be evaluated in future studies.

In conclusion, the effective restoration of GCase activity and correction of selected molecular and biochemical GD phenotypes highlight both the utility of the MLO platform for studying nGD and the therapeutic potential of this nanoparticle-based approach for treating nGD.

### AAV-mediated gene transfer in GD MLOs enhances GCase activity and mitigates GD phenotypes

AAV-based gene therapies have shown clinical safety and efficacy in neurodegenerative diseases ^59,60^. Trials for GD Type 1 (NCT05487599) and Type 2 (NCT04411654) using AAV9 have initiated. While animal models support AAV’s effectiveness in GD, its cellular impact on human or patient derived models remains unknown. To investigate the therapeutic effect of AAV gene therapy in human GD organoids, AAV9-GBA1 was injected at dose of 1.8 × 10^10^ vg per organoid at week 13 and evaluated 3 weeks after the injection (Figure 6A). In WT-75.1 MLOs, GCase activity was significantly increased in AAV9-GBA1-treated MLOs compared to untreated controls at week 15 (Figure 6B, left panel), suggesting effective transgene expression in normal organoids. AAV9-GBA1 restored GCase activity in GD2-1260 and GD2-10-257 MLOs to 47.8% and 37.7% of WT-75.1 levels, respectively (p < 0.001), compared to 6.0% and 8.8% in untreated controls (Figure 6B, middle and right panel). Immunoblot results further confirmed the restoration of GCase protein in AAV9-GBA1 treated GD MLOs (Figure 6B, bottom panel). Mass spectrometry analysis showed that AAV9-GBA1 normalized GluSph levels in GD2-1260 and GD2-10-257 MLOs, contrasting with elevated levels in untreated MLOs versus WT-75.1 controls (Figure 6C), indicating effective clearance of toxic lipid substrates in MLO by AAV gene therapy. Of note, LAMP1 levels were elevated after AAV9-GBA1 treatment, reflecting correction of lysosomal dysfunction (Figure 6D). However, the protein level of DA neuron marker (TH) was not significantly increased after AAV gene therapy compared to untreated group, suggesting longer treatment or earlier gene therapy intervention might be needed to recover dopaminergic neurons. Confocal imaging corroborated that AAV9-GBA1-treated GD2-1260 MLOs exhibited GFP signals and restored GCase expression in pan-neurons, DA neurons and astrocytes (Figure 6E).

**Figure 6.**
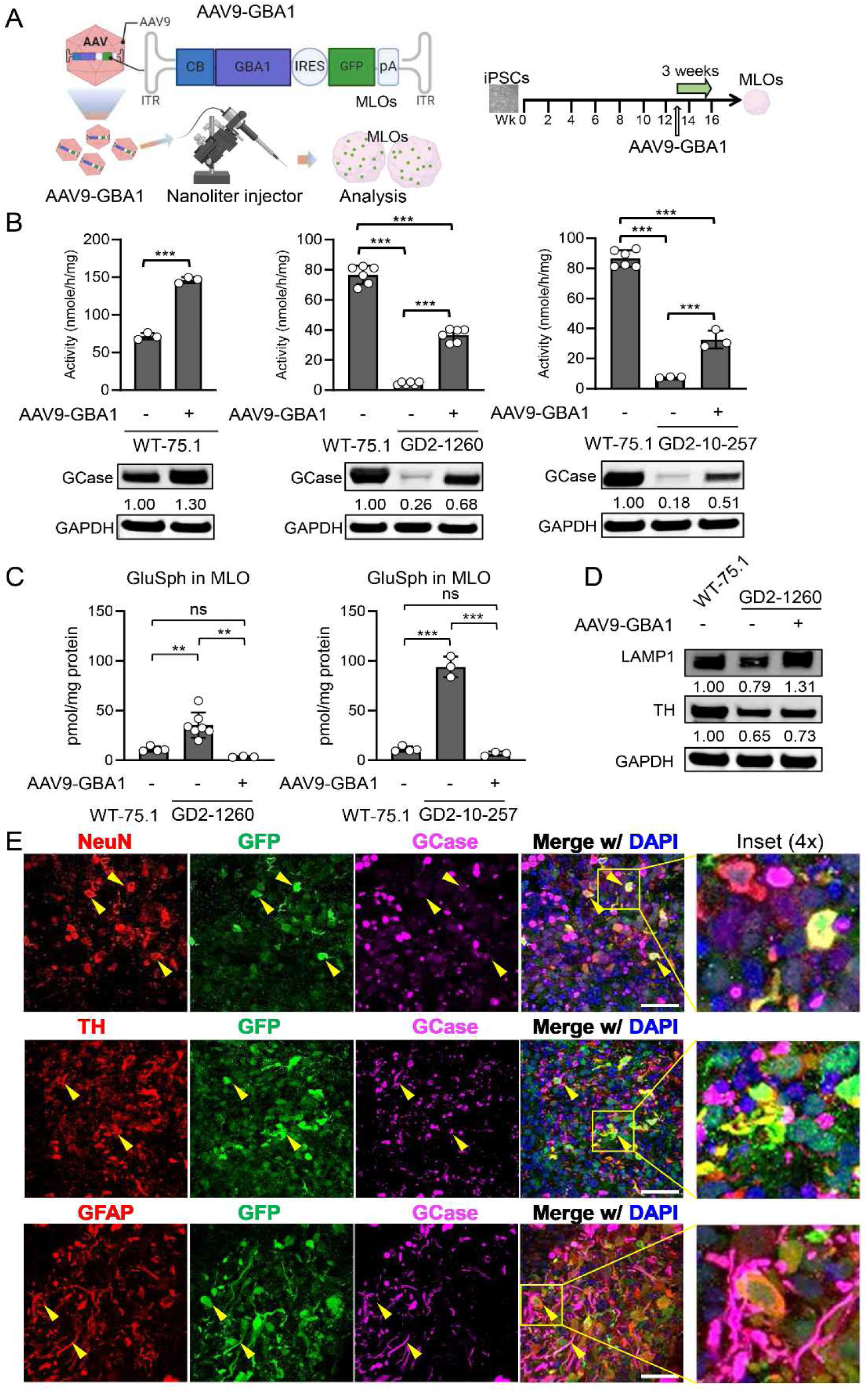
AAV9-GBA1 gene therapy mitigates disease phenotypes in GD MLOs. (A) Schematic illustration of AAV9-*GBA1* gene therapy delivery to MLOs using a nanoliter injector. AAV9 vectors carrying the GBA1 gene (AAV9-GBA1) are administered to Wk13 MLOs. The samples were analyzed after 3 weeks of treatment. (B) GCase activity in WT-75.1, GD2-1260, and GD2-10-257 MLOs and AAV9-GBA1 treated MLOs were measured by enzymatic assay. Data are presented as mean ± SEM (3 MLOs pooled, n = 3 to 6 per group). ***p < 0.001. (C) GluSph levels in AAV9-GBA1 treated GD and control MLOs were quantified by LC-MS/MS and normalized to total protein. Data are presented as mean ± SEM (3 MLOs pooled, n ≥ 3 per group). ***p < 0.001; ns, not significant. (D) Immunoblot analysis of LAMP1 and TH in WT-75.1 and in GD2-1260 MLOs untreated or treated with AAV9-GBA1. Protein samples were extracted from n=3 MLOs for each group. Protein levels are normalized to WT-75.1 untreated controls (set to 1.0). (E) Transgene expressions (yellow arrows and enlarged insert) in neurons (NeuN), DA neurons (TH) and astrocytes (GFAP) of AAV9-GBA1 treated GD2-1260 MLOs. Scale bar = 50 µm.

These results collectively demonstrate that AAV9-GBA1 gene therapy effectively corrected GCase deficiency, reduced GluSph accumulation, and ameliorates lysosomal pathology in GD MLOs, highlighting its therapeutic potential for nGD.

### Substrate Reduction Therapy reduces lipid accumulation and improves lysosomal function in GD MLOs

Current Substrate Reduction Therapy (SRT) in the standard clinical treatments for GD ^61,62^ has limited efficacy in addressing the neurological manifestations of nGD due to their inability to cross the blood-brain barrier efficiently. GZ-682452 (termed herein as GZ452) is an analogue of venglustat that is presently under clinical evaluation for treating nGD ^63^. To evaluate the human MLO as a preclinical model for SRT drug development, we assessed GZ452’s effects on MLO growth, lipid accumulation, midbrain markers, and lysosomal function.

GZ452 toxicity was first evaluated in WT-75.1 MLOs. Treatment at 0.3, 1, 2, 3 µM over 6 weeks modestly reduced MLO size at 1 µM, with the 2 µM and 3 µM doses significantly decreasing size versus untreated controls (Figure 7A), indicating high dose of GZ452 notably suppressed MLO growth. We next tested GZ452 at doses (0.01, 0.05, 0.3 µM) over 6 weeks in WT-75.1 MLOs. GZ452 dose-dependently reduced total GluCer levels (Figure 7B) and GluCer species, with the 0.3 µM dose showing the most pronounced effect (Figure 7C).

**Figure 7.**
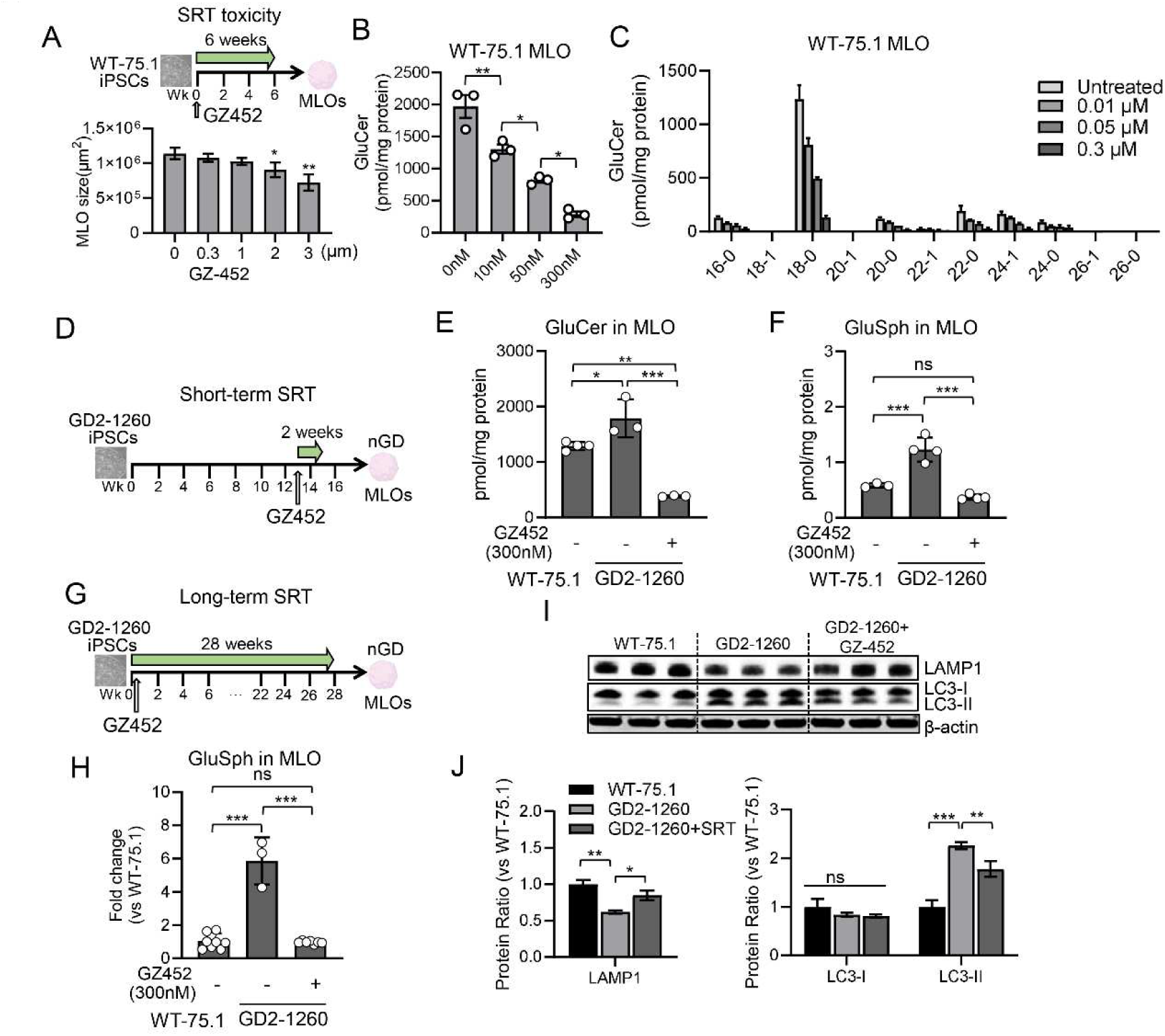
Substrate Reduction Therapy with GZ452 reduces lipid accumulation and improves autophagic and lysosomal abnormalities in GD MLOs. (A) Assessment of GZ452 tolerated dose in healthy MLO and the effect of GZ452 on organoid growth in WT-75.1 MLOs over 6 weeks. Three MLOs were pooled and n = 3 per concentration. (B, C) Total GluCer levels (B) and distribution of GluCer species (C) in WT-75.1 MLOs with various doses of GZ452 at Wk6. Three MLOs were pooled and n = 3 per concentration. (D) Schematic of the experimental timeline for short-term (2 weeks) GZ452 treatment of GD MLOs. (E, F) GluCer (E) and GluSph (F) levels in WT-75.1 and GD2-1260 MLOs at Wk15 under short-term GZ452 treatment. Data were normalized to protein mass. (G) Schematic of the experimental timeline for long-term (28 weeks) GZ452 treatment in GD MLOs. (H) GluSph levels in MLOs at Wk15 under long-term GZ452 treatment. Data were normalized to protein mass. (I) Immunoblot analysis of LAMP1 and LC3-I/II in SRT treated GD2-1260 MLOs for 28 weeks, with β-actin as loading control. Protein samples were extracted from n=3 MLOs for each group. (J) Quantification of LAMP1 and LC3-II/I in MLOs. Protein levels are normalized to WT-75.1 untreated controls (set to 1.0). All data presented as mean ± SEM (n = 3∼4). *p < 0.05; **p < 0.01; ns, not significant. One-way ANOVA with Tukey’s test.

Expression of midbrain markers *ASCL1*, *TH*, and *PLZF* in WT-75.1 MLOs was largely unaffected by GZ452 (0.3 µM) treatment started at week 6, with no significant changes in relative mRNA levels compared to untreated controls (Supplementary Figure 6). We therefore applied this optimal tolerated GZ452 dose of 0.3 µM (300nM) in short-term (2 weeks) and long-term (up to 28 weeks) treatment to test its therapeutic effects in GD MLOs. After two weeks of treatment starting in week 13, GZ452 significantly ameliorated GluCer and GluSph accumulation in GD2-1260 MLOs (Figure 7E, 7F). Long-term GZ452 treatment from day 2 to week 28 robustly cleared GluSph storage and normalized it to the WT level (Figure 7G, 7H). Immunoblot analysis showed that long-term GZ452 treatment improved lysosomal function, partially restoring LAMP1 levels (0.8 ± 0.1; p < 0.05) and reducing LC3-II by 21.4% (p < 0.01) (Figure 7I, 7J). These findings demonstrate that GZ452 effectively reduced lipid accumulation and improves lysosomal function in GD MLOs, supporting its therapeutic potential for ameliorating key disease phenotypes.

## Discussion

We developed MLOs from two GD Type 2 patient-derived hiPSCs, offering deepened understanding to disease pathogenesis and an advanced model system for drug discovery. To our knowledge, this is the first GD brain organoid model derived from patient cells. These GD MLOs harbor prevalent *GBA1*^L444P/P415R^ and *GBA1*^L444P/RecNcil^ mutations, replicated key nGD phenotypes, including reduced GCase activity, glycosphingolipids accumulation, lysosomal abnormalities, and impaired dopaminergic neuron differentiation (Supplementary Table 2). Emerging interventions, CRISPR/Cas9-mediated gene correction, SapC-DOPS-fGCase nanoparticle, and AAV9-GBA1 gene therapy, significantly restored GCase function and mitigate disease hallmarks in GD MLOs, supporting their use in preclinical studies (Supplementary Table 2). Additionally, dysregulated Wnt signaling and impaired dopaminergic neuron differentiation in GD MLOs provides novel insights into nGD pathogenesis and potential therapeutic targets. This research underscores the utility of patient-derived MLOs as a human-relevant platform for modeling disease complexity and testing drug responses to bridge preclinical and clinical research.

Our findings revealed nGD-specific phenotypes in MLOs, including reduced GCase activity, GluCer and GluSph accumulation, and altered mTOR and autophagosome-lysosomal pathways, closely reflecting patient neuropathology ^4,55^. Unlike *Gba1* knockout animal models, which lack patient-specific genetic and epigenetic diversity, patient-derived MLOs retain heterozygous or compound heterozygous background, enabling study of disease mechanisms and modifier genes. Partial rescue in isogenic iso-GD2-1260 MLOs, where mutation correction alleviated but did not fully resolve abnormalities, highlights the model’s value in capturing nGD pathology and guiding personalized precision therapeutic strategies. This model offers a more advanced and physiologically relevant platform than traditional gene knockout or transgenic models, better capturing the complexity of nGD pathology.

Transcriptomic analysis of GD MLOs uncovered dysregulated Wnt signaling, a critical pathway in early brain development, highlighting a mechanistic insight into nGD pathogenesis. Aligning with Awad et al., who reported that downregulated Wnt/β-catenin signaling impairs DA neuron differentiation in nGD hiPSC-derived neuronal progenitors ^30^, our MLOs exhibited reduced FOXA2+ DA progenitors and TH+ mature DA neurons, along with decreased dopamine release in culture medium. Transcriptome analysis of the GD MLOs identified 1,429 differentially expressed genes enriched in Wnt signaling, axon guidance and neuron differentiation pathways, indicating that GCase deficiency disrupts midbrain patterning and neuronal specification. This finding is further supported by reduced expression of midbrain markers (e.g., FOXA2, TH), reduced dopamine release from DA neurons, and skewed forebrain/midbrain patterning genes (e.g., FOXP1, FOXG1) in GD MLOs. Compared to *GBA1* KO iPSC-derived midbrain-like organoids, which showed only a marginal reduction in dopaminergic neuron differentiation and no significant dysregulation of FOXG1 expression ^24^, our MLO model exhibits a distinct and more disease-relevant phenotype. These results suggest that aberrant Wnt signaling contributes to impaired dopaminergic neuron development and may drive the disease pathogenesis in nGD. Thus, targeting Wnt-related pathways could offer a therapeutic strategy to address early developmental defects in nGD. Comparative analysis with prior transcriptomic data from nGD mouse midbrain showed consistent dysregulation in axon guidance, synaptic signaling, lipid metabolism, and nervous system development etc. (Supplementary Table 3) ^13^, supporting the fidelity of our human MLO model. Moreover, midbrain organoids at 35 days of development closely correspond to the human embryonic midbrain at approximately 9 weeks, while 70-day-old organoids align more closely with the 10-week stage ^64^. These correlations highlight the importance of initiating treatment during the fetal stage, which may be critical for preventing the disease.

Isogenic MLOs generated by correcting the *GBA1* L444P mutation in GD2-1260 hiPSCs using CRISPR/Cas9 revealed the key genetic mechanisms in nGD. The resulting WT/P415R genotype in iso-GD2-1260 partially rescued nGD phenotypes. GCase activity restored to nearly 50% of WT-75.1 levels, with normalized GluSph levels and improved TH expression. However, partial recovery of FOXA2 (53% of WT-75.1), along with persistent LC3 and LAMP1 abnormalities, may contribute to remaining pathology. These findings imply that the retained P415R mutation in this isogenic line may contribute to residual lysosomal dysfunction, potentially exerting a negative effect, as it has been reported to be associated with altered GCase stability and activity in previous studies ^65,66^. The partial rescue in iso-GD2-1260 MLOs suggests that the P415R mutation may affect disease severity and therapeutic response, deserving further investigation into the specific role of P415R mutation in GD and assessing whether dual mutation correction is required for full phenotypic normalization.

Evaluation of GZ452, a CNS-accessible SRT drug ^67^, in GD2-1260 MLOs showed significant reduction in GluCer and GluSph levels at a well-tolerated dose, alongside partial restoration of lysosomal and autophagy markers (LAMP1, LC3-II). Unlike higher doses that affected organoid growth, 0.3 µM preserved MLO size and expression of midbrain markers (ASCL1, TH, PLZF) involved in organoid differentiation and growth. These findings underscore the value of the MLOs for assessing drug safety (e.g., organoid size as toxicity indicator) and efficacy in a human-specific, midbrain-relevant model before clinical studies. Our study demonstrated positive therapeutic outcomes using AAV9-GBA1 gene therapy and SapC-DOPS nanoparticle-mediated GCase delivery in GD2-1260 and GD2-10-257 MLOs. AAV9-GBA1 restored GCase activity, while SapC-DOPS-fGCase elevated GCase activity to WT levels, both approaches effectively reduced GluSph. Based on prior studies showing no region-specific effects in GD patients and mice, these therapies may also be effective in other brain organoid types, such as COs ^4,41^. The novelty of employing MLOs lies in their ability to concurrently test these diverse therapeutic modalities: SRT, gene therapy, and nanoparticle-based enzyme delivery within a patient-derived system that replicates nGD pathology, offering the unique patient-MLO system to assess complex cellular responses that directly reflect patient specific conditions.

Unlike animal models, MLOs provide a controlled environment to evaluate dose-dependent effects and cellular responses, enabling the identification of optimal therapeutic windows while minimizing the risk of adverse outcomes in patients with nGD. Moreover, FDA has increasingly encouraged the use of organoid systems in drug development, recognizing their potential to improve preclinical testing and reduce reliance on animal models. Therefore, the adaptability of MLOs to test advanced interventions, coupled with their alignment with regulatory trends favoring human-relevant models, positions them as a great tool for accelerating the translation of innovative treatments from preclinical research to clinical application.

There are limitations in our current MLOs, such as lacking vascular system and microglia. The absence of microglia and vasculature may contribute to incomplete phenotyping and phenotypic rescue observed in our therapeutic experiments. Without vascularization, cells in the core may experience hypoxic and nutrient deprivation, leading cell death or necrosis in deeper layers in long-term culture. This limitation may partially explain the incomplete restoration of dopaminergic markers (TH, FOXA2) and partial recovery of LAMP1 following SapC-DOPS-fGCase and AAV9-GBA1 treatments. Additionally, the lack of microglia limits molding of neuroinflammatory, which contributes to the clearance of toxic lipid substrates and mediating neuronal damage in GD. Elevated GluSph and impaired dopaminergic differentiation observed in GD MLOs may not fully reflect inflammatory process. Dysregulated Wnt signaling and lysosomal and autophagic markers (LAMP1, LC3-II/I) could be amplified by microglial activation *in vivo*. To improve model fidelity, future studies would incorporate vascularization (e.g., endothelial co-culture or microfluidics) to enhance oxygen and nutrient distribution and integrate microglia via directed differentiation ^68^ or co-culture with hiPSC-derived microglia ^69^ to enhance the relevance of MLOs for long-term nGD research and drug testing.

In conclusion, patient-derived MLOs offer an advanced, human-relevant platform for elucidating nGD pathogenesis and evaluating therapies, overcoming limitations of traditional models. Dysregulated Wnt signaling and impaired dopaminergic neuron differentiation, along with partial rescue of phenotypes through gene correction, highlight the complex genetic and developmental underpinnings of nGD. The efficacy assessment of SapC-DOPS-fGCase, AAV9-GBA1 and SRT-GZ452 therapies underscores the targeted interventions and safety evaluation of these approaches in patient-derived brain organoid models. While limitations such as lack of vascularization and microglia constrain modeling of neuroinflammation, these challenges can be addressed as discussed above and using advanced culturing techniques. Collectively, our findings lay the groundwork for developing patient-specific preclinical models to support personalized therapeutic strategies, while also advocating for the continued refinement of MLO systems to accelerate the discovery of effective treatments for nGD.

## Acknowledgments

This work was supported by the Cincinnati Children’s Pluripotent Stem Cell Facility (RRID: SCR_022634), Clinical and Biomedical Mass Spectrometry Facility (RRID: SCR_022638) and Transgenic Animal and Genome Editing Facility (RRID: SCR_022642). Authors acknowledge Freeline Therapeutics (now Spur Therapeutics) for providing the fGCase under a Material Transfer Agreement.

## Additional information

## Author Contributions

Y.L. designed experiments, cultured hiPSCs and MLOs, performed experiments, analyzed data, and wrote the manuscript. B.L. measured GCase enzyme activity in hiPSCs and MLOs. V.F. carried out lipid extract from hiPSCs and MLOs, W.Z., X.Z., R.L.B. and K.DR.S performed quantification of GluCer/GluSph by LC-MS/MS and data analysis. S.A. assisted with immunofluorescence staining and immunoblot assays. C.N.M and YC.H made isogenic hiPSC cells and conceived experimental designs and edited the manuscript. J.T. conceived experimental designs and edited the manuscript. J.E.H. generated cerebral organoid and edited manuscript. A.K. and X.Q. provided SapC-DOPS nanoparticles and guidance for the enzyme uptake assay. R.A.F provided an nGD hiPSC cell line and edited the manuscript. Y.S. conceived experimental designs, supervised the project, wrote the manuscript, and provided funding to the study.

## Ethics

This work has been reviewed and approved by the Cincinnati Children’s Hospital Medical Center (CCHMC) and University of Cincinnati (UC) Embryonic Stem Cell Research Oversight (ESCRO) committee under ESCRO protocol #EIPHG3114_R3. Primary fibroblast (GM01260) for generating GD2-1260 iPSC was obtained from the Coriell Institute for Medical Research under Material Transfer Agreement. WT-75.1 iPSC was generated using fibroblasts cultured from discarded circumcision tissue obtained from a healthy donor (provided by Dr. Susanne Wells, Division of Hematology/Oncology, CCHMC; institutional review board no. 02-9-29X) at CCHMC. The procedures used to generate the GD2-1260 iPSC and WT-75.1 iPSC lines at CCHMC have been reviewed and approved by CCHMC/UC ESCRO committee. Skin biopsies to generate GD2-10-257 iPSCs were obtained from NIH with informed consent and were approved by the Institutional Review Board (IRB). The procedures used to generate the GD-2-10-257 nGD iPSC line at the University of Maryland Baltimore (UMB) have been reviewed and approved by the UMB IRB and ESCRO committees.

## Additional files

## Supplementary files

**Supplementary Fig. 1.** Correct L444P mutation in nGD iPSC by CRISPR-Cas9. (A) Single stranded oligonucleotide design. Silent mutations in upper case. Phosphorothioate modified bases (*). (B) Genomic sequence of mutant, wild type and corrected GBA1. Mutated codon highlighted in red. T insertion for point mutation correction highlighted in yellow. PAM highlighted in grey. sgRNA target sequence underlined. (C) DNA electrophoresis gels showing genome editing and clone screening for iso-GD2-1260 (clone #9). (D) Karyotyping of GD2-1260 and CRISPR/Cas9 corrected iso-GD2-1260 hiPSCs. Normal karyotype was observed in both hiPSC lines. (E) Relative mRNA expression of genes (OCT4, NANOG and SOX2) required for generating and maintaining hiPSCs pluripotency by quantitative RT-PCR. ns, not significant by One-way ANOVA analysis.

**Supplementary Fig. 2.** Representative UHPLC-MS/MS chromatograms of GluSph and GluCer species in WT-75.1 and GD2-1260 MLOs. Tissues from Wk 28 MLOs were tested.

**Supplementary Fig. 3**. Neural rosette formation during MLO maturation were not affected by GBA1 mutation. (A) Cryosections of Wk 6 MLOs derived from WT-75.1, GD2-1260 and isogenic control iso-GD2-1260 hiPSC cells. Sections were stained with antibodies against Ki67 and Sox2 and nuclei were costained with DAPI. Scale bar, 100 µm. (B) Quantification of the Sox2+, Ki67+ and Sox2+/Ki67+ cells (mean ± SEM; n = 3). ns, not significant by One-way ANOVA analysis.

**Supplementary Fig. 4**. Restoration of GCase Expression in dopaminergic neuron and astrocytes in SapC-DOPS-fGCase treated nGD MLOs. (A, B) Representative confocal images of GD2-1260 (A) and GD2-10-257 MLOs (B) treated with SapC-DOPS-fGCase for 2 weeks, immunostained for GCase (green) and dopaminergic neuron (TH, red) with DAPI (blue) labeling nuclei. (C, D) Representative immunostaining images for GCase (green) and astrocytes (GFAP, red) in GD2-1260 (C) and GD2-10-257 MLOs (D) treated with SapC-DOPS-fGCase. Scale bar, 50 µm. Yellow arrows indicate colocalized GCase in TH+ cells. (E) Immunoblot of TH in Wk 16 and Wk 28 GD2-10-257 MLOs.

**Supplementary Fig. 5**. Restoration of GCase expression in lysosomal and autophagosomal compartments in SapC-DOPS-fGCase treated nGD MLOs. (A, B) Representative confocal images of GD2-1260 (A) and GD2-10-257 MLOs (B) treated with SapC-DOPS-fGCase for 2 weeks, immunostained for GCase (green) and lysosomal marker LAMP1 (red) with DAPI (blue) labeling nuclei. Yellow arrows indicate colocalized GCase in LAMP1+ compartments. Scale bar, 50 µm. (C, D) Representative confocal images of GD2-1260 (C) and GD2-10-257 MLOs (D) treated with SapC-DOPS-fGCase for 2 weeks, immunostained for GCase (green) and autophagosomal marker LC3B (red) with DAPI (blue) labeling nuclei. Yellow arrows indicate colocalized GCase in LC3B+ compartments. Scale bar, 50 µm.

**Supplementary Fig. 6**. Influence of SRT drug GZ452 on DA neuron differentiation in WT-75.1 MLOs. Relative mRNA expression of midbrain markers ASCL1, TH, and PLZF in WT-75.1 at Wk6 in untreated or treated MLOs with indicated concentrations of GZ452. Relative gene expression is normalized to untreated control WT-75.1 MLOs (set to 1.0).

**Supplementary Table 1**. Key Resources

**Supplementary Table 2**. Summary of therapeutic modalities on nGD MLOs

**Supplementary Table 3**. Key Dysregulated Pathways

## Data availability

RNA-seq data generated in this study were deposited in Gene Expression Omnibus (GEO) at National Institutes of Health (NIH) with accession number GSE303993.

