## supplemental file for "Patient-Specific Midbrain Organoids with CRISPR Correction Recapitulate Neuronopathic Gaucher Disease Phenotypes and Enable Evaluation of Novel Therapies"

**Supplementary Figures**

**Supplementary Fig. 1.** Correct L444P mutation in nGD hiPSC by CRISPR-Cas9.

**Supplementary Fig. 2.** Representative UHPLC-MS/MS chromatograms.

**Supplementary Fig. 3.** Neural rosette formation during MLO maturation was not affected by *GBA1* mutation.

**Supplementary Fig. 4.** Restoration of GCase Expression in dopaminergic neuron and astrocytes in SapC-DOPS-fGCase-treated nGD MLOs.

**Supplementary Fig. 5.** Restoration of GCase expression in lysosomal and autophagosomal compartments in SapC-DOPS-fGCase-treated nGD MLOs.

**Supplementary Fig. 6.** Influence of SRT drug GZ452 on DA neuron differentiation in WT-75.1 MLOs.

**Supplementary Table 1. Key Resources**

**Supplementary Table 2. Summary of therapeutic modalities on nGD MLOs**

**Supplementary Table 3. Dysregulated pathways in nGD models**

### Supp. Fig. 1

A Single stranded oligonucleotide design

t*c*t*tcagcccacttcccagacctcaccattgccctcaccggtttagcacgaccacaacagcagagccatcgggatgcatcagtgccacGgcAt
ccAgAtcgttcttctgactggcaaccagc*c*c*c

B Genomic sequence

```
L444P Mut GBA1:    gtt gcc agt cag aag aac gac ccg gac gca gtg gca ctg atg
Wild type GBA1:    gtt gcc agt cag aag aac gac ctg gac gca gtg gca ctg atg
Corrected GBA1:    gtt gcc agt cag aag aac gaT ctg gaT gcC gtg gca ctg atg
```

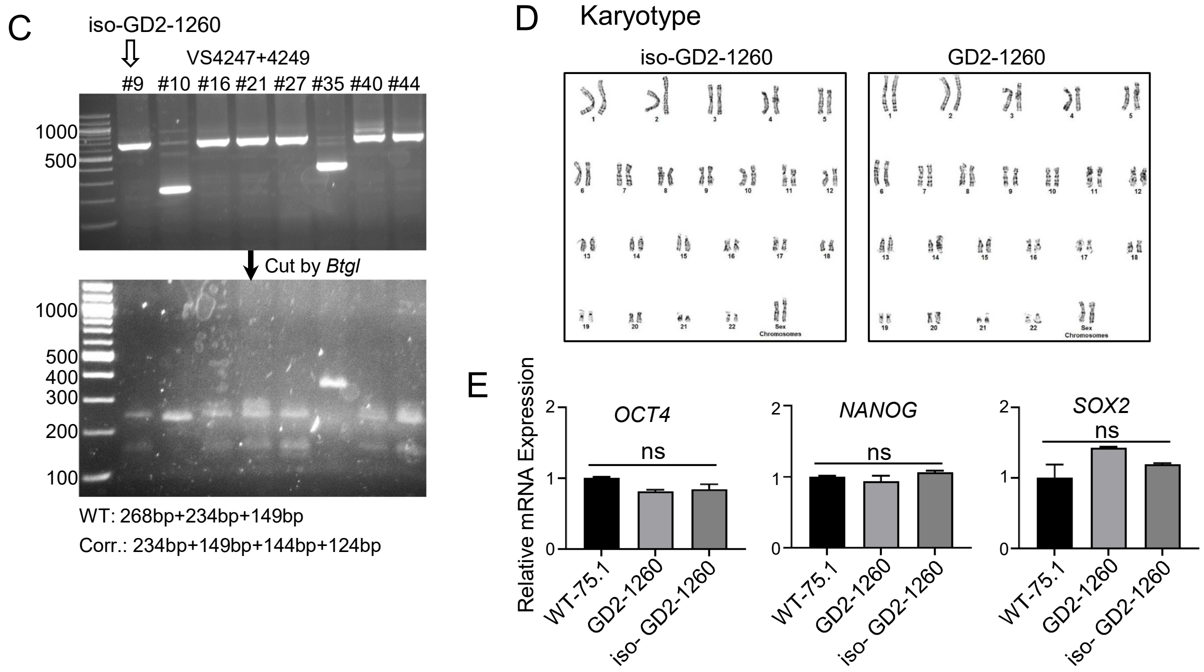

**Supplementary Fig. 1 Correct L444P mutation in nGD iPSC by CRISPR-Cas9.** (A) Single stranded oligonucleotide design. Silent mutations in upper case. Phosphorothioate modified bases (*). (B) Genomic sequence of mutant, wild type and corrected GBA1. Mutated codon highlighted in red. T insertion for point mutation correction highlighted in yellow. PAM highlighted in grey. sgRNA target sequence underlined. (C) DNA electrophoresis gels showing genome editing and clone screening for iso-GD2-1260 (clone #9). (D) Karyotyping of GD2-1260 and CRISPR/Cas9 corrected iso-GD2-1260 hiPSCs. Normal karyotype was observed in both hiPSC lines. (E) Relative mRNA expression of genes (OCT4, NANOG and SOX2) required for generating and maintaining hiPSCs pluripotency by quantitative RT-PCR. ns, not significant by One-way ANOVA analysis.

**Supp. Fig. 2**

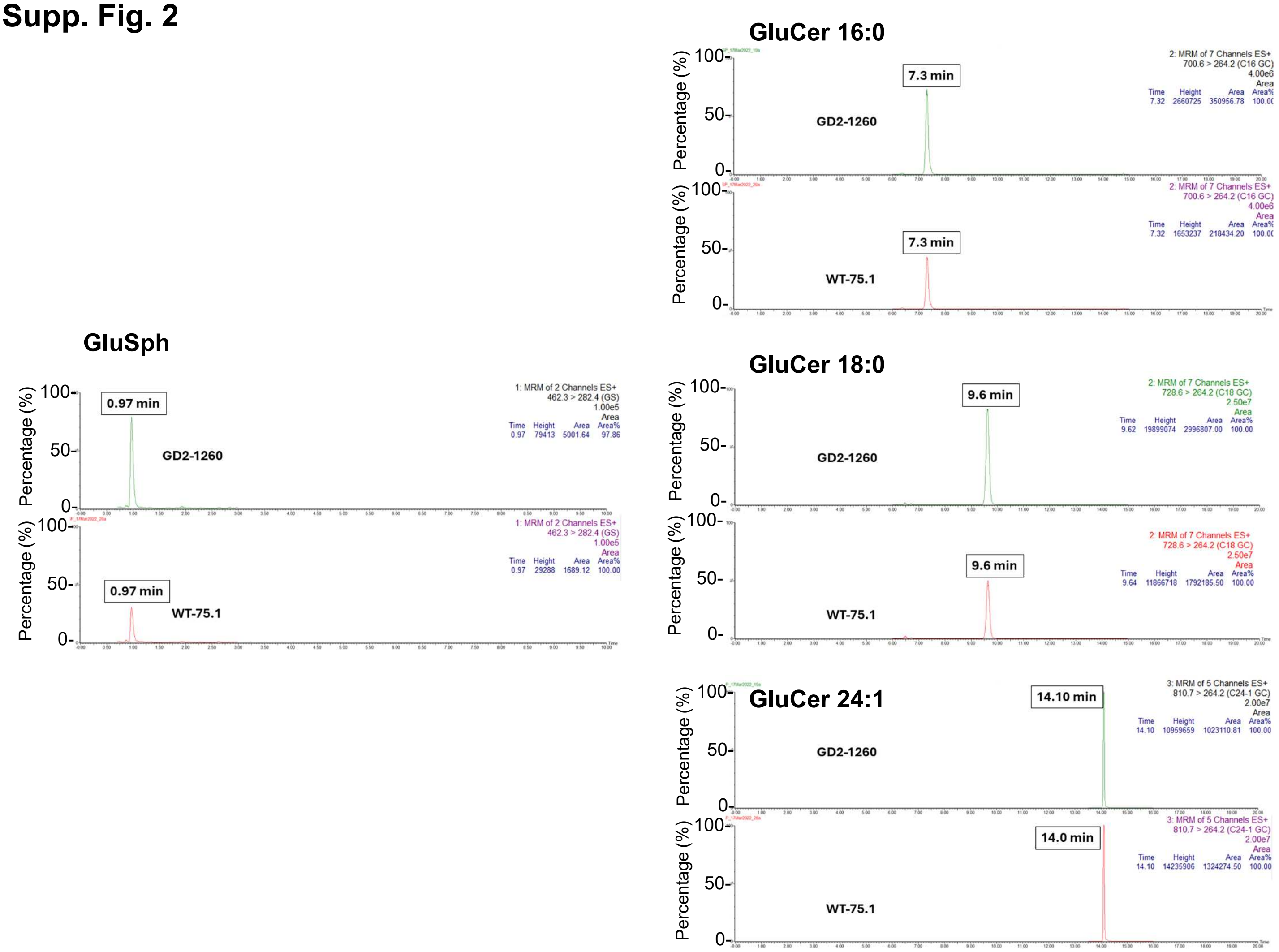

**Supplementary Fig. 2.** Representative UHPLC-MS/MS chromatograms of GluSph and GluCer species in WT-75.1 and GD2-1260 MLOs. Tissues from Wk 28 MLOs were tested.

#### Supp. Fig. 3

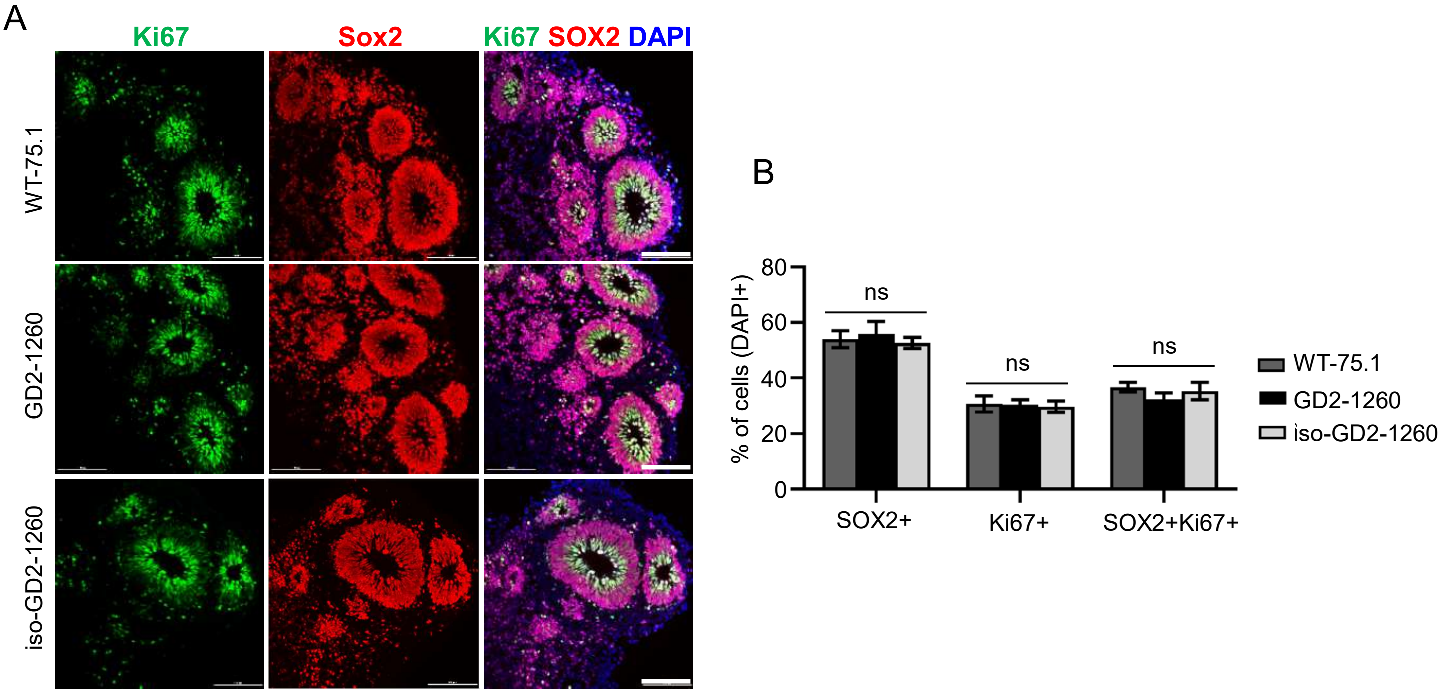

**Supplementary Fig. 3. Neural rosette formation during MLO maturation were not affected by *GBA1* mutation.** (A) Cryosections of Wk 6 MLOs derived from WT-75.1, GD2-1260 and isogenic control iso-GD2-1260 hiPSC cells. Sections were stained with antibodies against Ki67 and Sox2 and nuclei were costained with DAPI. Scale bar, 100 µm. (B) Quantification of the Sox2+, Ki67+ and Sox2+/Ki67+ cells (mean ± SEM; n = 3). ns, not significant by One-way ANOVA analysis.

#### Supp. Fig. 4

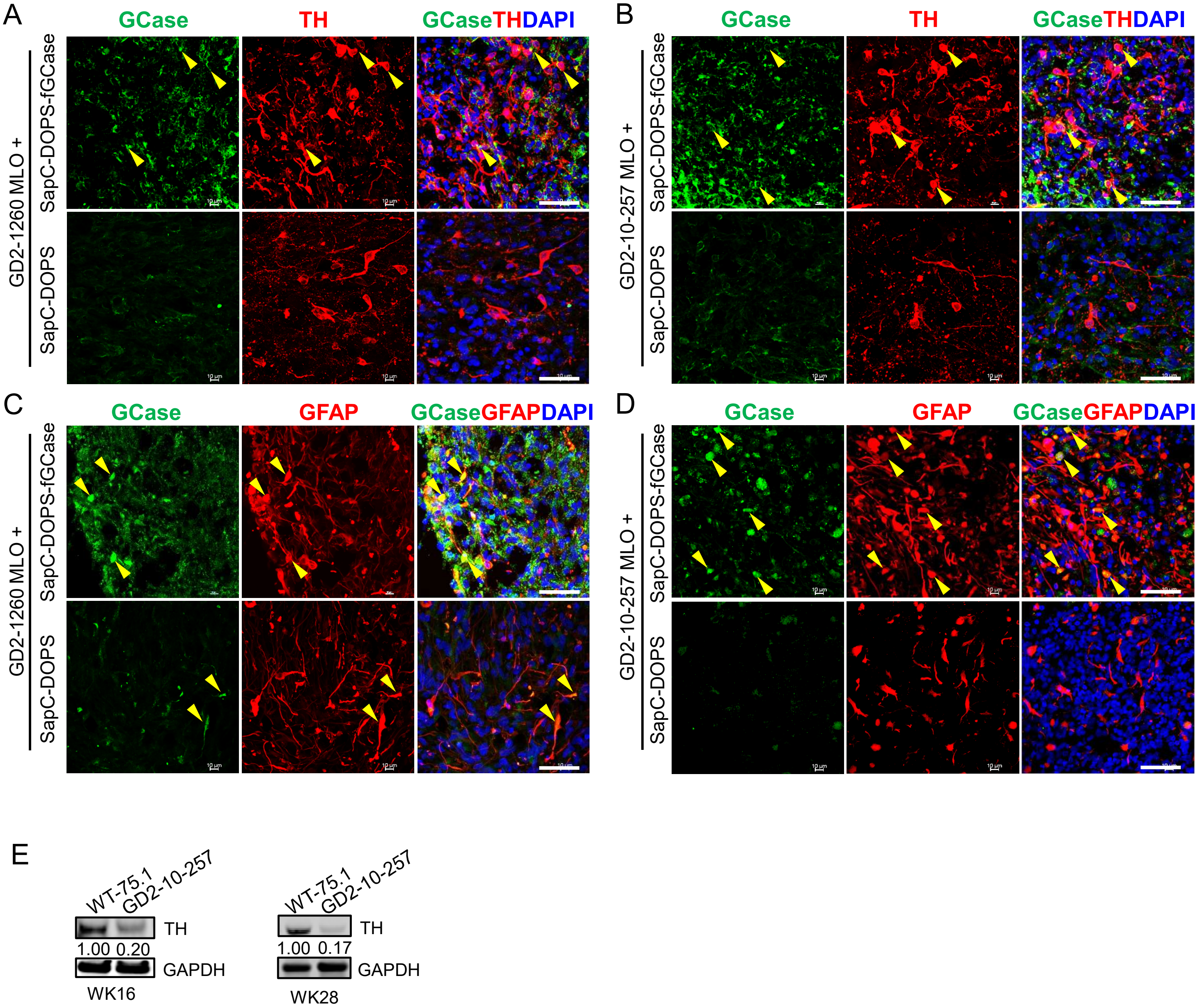

**Supplementary Fig. 4. Restoration of GCase Expression in dopaminergic neuron and astrocytes in SapC-DOPS-fGCase-treated nGD MLOs.** (A, B) Representative confocal images of GD2-1260 (A) and GD2-10-257 MLOs (B) treated with SapC-DOPS-fGCase for 2 weeks, immunostained for GCase (green) and dopaminergic neuron (TH, red) with DAPI (blue) labeling nuclei. (C, D) Representative immunostaining images for GCase (green) and astrocytes (GFAP, red) in GD2-1260 (C) and GD2-10-257 MLOs (D) treated with SapC-DOPS-fGCase. Scale bar, 50 µm. Yellow arrows indicate colocalized GCase in TH+ cells. (E) Immunoblot of TH in Wk 16 and Wk 28 GD2-10-257 MLOs.

#### Supp. Fig. 5

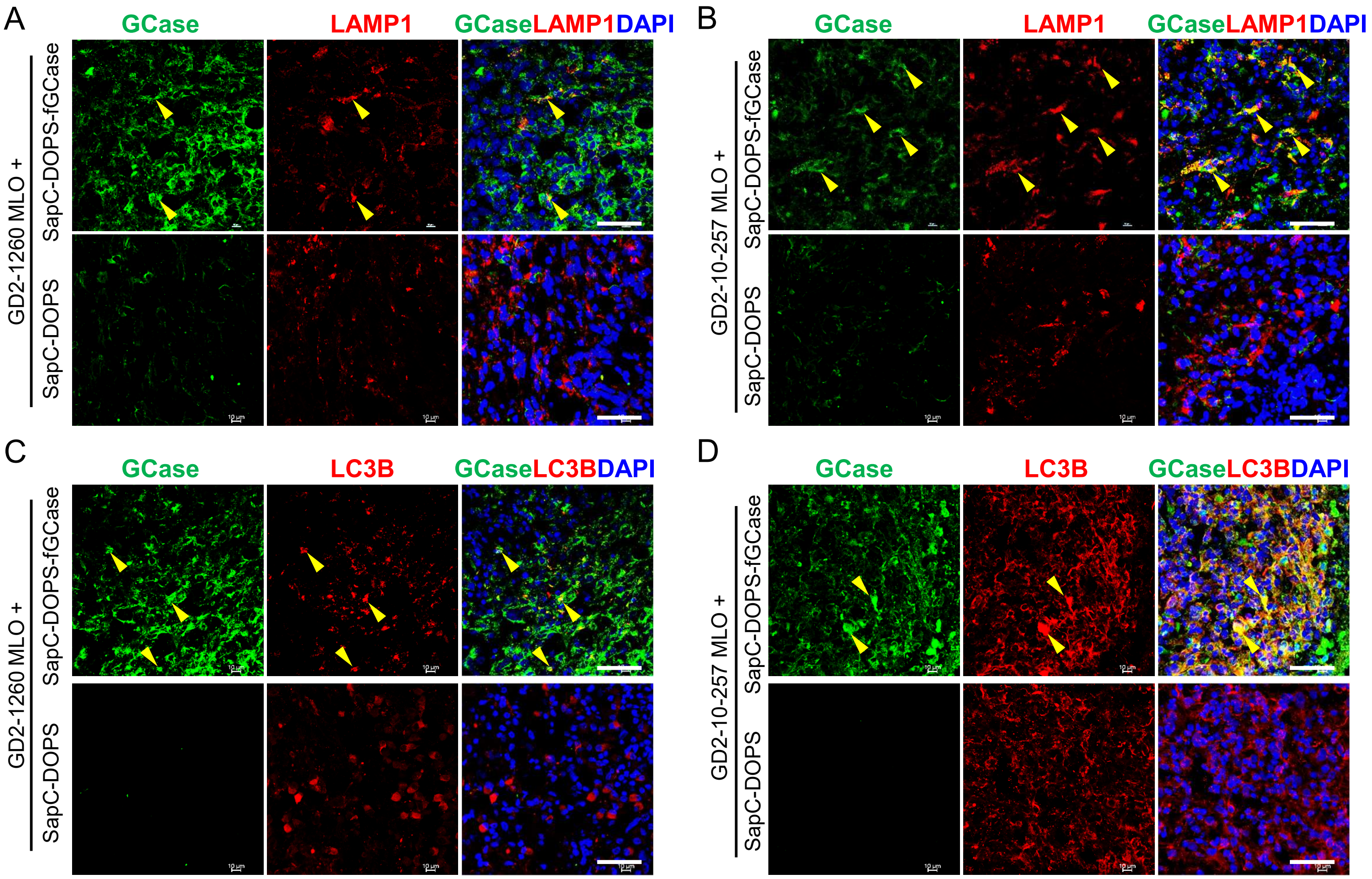

**Supplementary Fig. 5. Restoration of GCase expression in lysosomal and autophagosomal compartments in SapC-DOPS-fGCase-treated nGD MLOs.** (A, B) Representative confocal images of GD2-1260 (A) and GD2-10-257 MLOs (B) treated with SapC-DOPS-fGCase for 2 weeks, immunostained for GCase (green) and lysosomal marker LAMP1 (red) with DAPI (blue) labeling nuclei. Yellow arrows indicate colocalized GCase in LAMP1+ compartments. Scale bar, 50 µm. (C, D) Representative confocal images of GD2-1260 (C) and GD2-10-257 MLOs (D) treated with SapC-DOPS-fGCase for 2 weeks, immunostained for GCase (green) and autophagosomal marker LC3B (red) with DAPI (blue) labeling nuclei. Yellow arrows indicate colocalized GCase in LC3B+ compartments. Scale bar, 50 µm.

**Supp. Fig. 6**

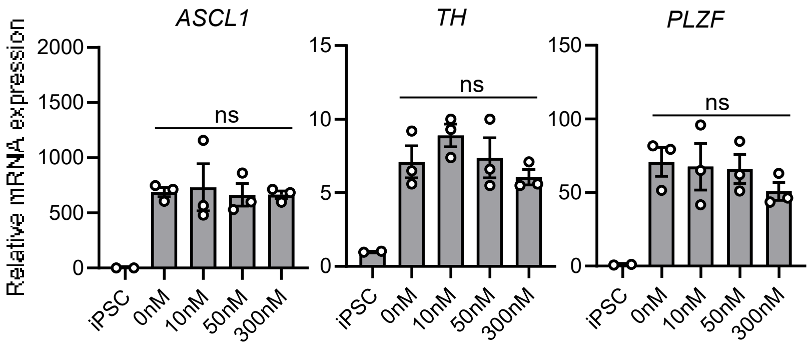

**Supplementary Fig. 6. Influence of SRT drug GZ452 on DA neuron differentiation in WT-75.1 MLOs.** Relative mRNA expression of midbrain markers *ASCL1*, *TH*, and *PLZF* in WT-75.1 at Wk6 in untreated or treated MLOs with indicated concentrations of GZ452. Relative gene expression is normalized to untreated control WT-75.1 MLOs (set to 1.0).

#### Supplementary Table 1. Key Resources

| REAGENTS and SUPPLIES | SOURCE | IDENTIFIER |
|---|---|---|
| ***Antibodies*** | | |
| Tuj1 | BioLegend | Cat#801201; RRID:AB_2313773 |
| NeuN | Millipore | Cat#MAB377; RRID:AB_2298772 |
| FOXA2 | Cell Signaling Technology | Cat#8186S; RRID:AB_10891055 |
| GFAP | STEMCELL Technologies | Cat#60048.1; RRID:AB_3095092 |
| TH | Millipore | Cat#AB152; RRID:AB_390204 |
| TH | Cell Signaling Technology | Cat#45648S; RRID: AB_3677640 |
| 4e-bp1 | Cell Signaling Technology | Cat#9452S; RRID:AB_331692 |
| Phospho-4E-BP1 (Thr37/46) | Cell Signaling Technology | Cat#2855S; RRID:AB_560835 |
| β-Actin | Invitrogen | Cat#MA5-15739; RRID:AB_10979409 |
| Cathepsin D | Novusbio | Cat#NBP2-67477; RRID:AB_3095093 |
| FOXG1 | Abcam | Cat#ab18259; RRID:AB_732415 |
| FOXP1 | Millipore | Cat#MAB45341; RRID:AB_3658314 |
| GAPDH | Millipore | Cat#MAB374; RRID:AB_3658314 |
| GFP | Abcam | Cat#ab13970; RRID:AB_300798 |
| GFP | Invitrogen | Cat#A11120; RRID:AB_221568 |
| hGCase (NY#10, pAb) | Made in lab | Made in lab, NY#10; RRID: AB_3677641 |
| SOX2 | Cell Signaling Technology | Cat#23064S; RRID:AB_2714146 |
| Ki67 | Cell Signaling Technology | Cat#9449S; RRID:AB_2797703 |
| Lamp1 (CD107a) | Bioss | Cat#bsm-51301M; RRID: AB_3677642 |
| LC3B | Novusbio | Cat#NB100-2220; RRID:AB_10003146 |
| MAP2 | Cell Signaling Technology | Cat#4542S; RRID:AB_10693782 |
| PAX6 | Covance | Cat#14811801; RRID:AB_2315064 |
| S6 Ribosomal Protein | Cell Signaling Technology | Cat#2217S; RRID:AB_331355 |
| Phospho-S6 Ribosomal Protein (Ser235/236) | Cell Signaling Technology | Cat#4856S; RRID:AB_2181037 |
| ***Chemicals, Peptides, and Recombinant Proteins*** | | |
| Vitronectin | ThermoFisher | Cat#A14700 |
| Y-27632; ROCK inhibitor | Tocris Bioscience | Cat#1254 |
| CEPT | Bio-Techne | Cat#7991 |
| FGF-Basic (FGF-b, human) | ThermoFisher | Cat#PHG0264 |
| Dorsomorphin | Millipore | Cat#171261-1MG |
| A83-01 | STEMCELL Technologies | Cat#72024 |
| CHIR99021 | STEMCELL Technologies | Cat#72052 |
| IWP2 | STEMCELL Technologies | Cat#72122 |
| FGF8 | Thermofisher | Cat#100-25A-100UG |
| SAG | STEMCELL Technologies | Cat#73412 |
| Laminin | STEMCELL Technologies | Cat#77003 |
| BDNF | STEMCELL Technologies | Cat#78005.1 |

| | | |
|---|---|---|
| GDNF | STEMCELL Technologies | Cat#78058.1 |
| Ascorbic acid | Peprotech | Cat#5088177 |
| db-cAMP | Sigma aldrich | Cat#D0627-250MG |
| Growth Factor Reduced (GFR) Matrigel | Corning | Cat#354230 |
| ***Critical Commercial Assays*** | | |
| CEPT | Bio-Techne | Cat#7991 |
| Dopamine ELISA Kit | Abnova | Cat#KA3838 |
| ***Experimental Models: iPSC Lines*** | | |
| WT-75.1 | | CVCL_C1UB (PMCID=PMC9391520); https://www.cellosaurus.org/CVCL_C1UB |
| GD2-1260 | | https://pmc.ncbi.nlm.nih.gov/articles/PMC4378893/ |
| GD2-10-257 | | https://www.sciencedirect.com/science/article/pii/S2213671117304848; |
| ***Oligonucleotides*** | ***5' to 3'*** | |
| *NANOG FP* | TGCAACCTGAAGACGTGTGA | DOI: 10.1002/stem.3163 Stem Cells. 2020;38:727 – 740. |
| *NANOG RP* | CTATGAGGGATGGGAGGA | DOI: 10.1002/stem.3163 Stem Cells. 2020;38:727 – 740. |
| *OCT4 FP* | GACAGGGGGAGGGGAGGAGCTAGG | DOI: 10.1002/stem.3163 Stem Cells. 2020;38:727 – 740. |
| *OCT4 RP* | CTTCCCTCCAACCAGTTGCCCCAAAC | DOI: 10.1002/stem.3163 Stem Cells. 2020;38:727 – 740. |
| *PLZF FP* | TCCCGCCCGACTGGAGGATA | DOI: 10.1002/stem.3163 Stem Cells. 2020;38:727 – 740. |
| *PLZF RP* | TTCTTTCCTGGCTCCCCGCTC | DOI: 10.1002/stem.3163 Stem Cells. 2020;38:727 – 740. |
| *FOXG1 FP* | GCGGGCCAGACCAGTTACTT | DOI: 10.1002/stem.3163 Stem Cells. 2020;38:727 – 740. |
| *FOXG1 RP* | CCCAGACAGTCCCGTCGTAA | DOI: 10.1002/stem.3163 Stem Cells. 2020;38:727 – 740. |
| *TH FP* | CTGAGATTCGGGCCTTCGAC | DOI: 10.1002/stem.3163 Stem Cells. 2020;38:727 – 740. |
| *TH RP* | TGCACCTAGCCAATGGCACT | DOI: 10.1002/stem.3163 Stem Cells. 2020;38:727 – 740. |
| *ASCL1 FP* | GGTGATCGCACAACCTGCAT | DOI: 10.1002/stem.3163 Stem Cells. 2020;38:727 – 740. |
| *ASCL1 RP* | GTTCTGAGCGCTTCCCGTTT | DOI: 10.1002/stem.3163 Stem Cells. 2020;38:727 – 740. |
| *SOX2 FP* | AGACTGCACATGAGCCAGCA | This study. |
| *SOX2 RP* | CGTCTCCAGCCAGCTTCAAC | This study. |
| *FOXA2 FP* | TACGACGACATGTTCATGGAGC | This study. |
| *FOXA2 RP* | TATGCTGGGAGCGGTGAAGA | This study. |
| *GFAP FP* | CTGTTGCCAGAGATG GAGGTT | https://doi.org/10.1038/s41467-020-19264-0 |
| *GFAP RP* | TCATCGCTCAGGAGGTCCTT | https://doi.org/10.1038/s41467-020-19264-0 |
| *S100B FP* | GGAGACGGCGAATGTGACTT | DOI: https://doi.org/10.7554/eLife.52904 |
| *S100B RP* | GAACTCGTGGCAGGCAGTAGTAA | DOI: https://doi.org/10.7554/eLife.52904 |
| *GLAST FP* | CCAGCAGGGAGTCCGTAAAC | DOI: https://doi.org/10.7554/eLife.52904 |

| | | |
|---|---|---|
| *GLAST RP* | GCAGCACAAAAGCATTCCGA | DOI: https://doi.org/10.7554/eLife.52904 |
| *GAPDH FP* | TTGAGGTCAATGAAGGGGTC | This study. |
| *GAPDH RP* | GAGGTGAAGGTCGGAGTCA | This study. |
| *ACTB FP* | CTGGCACCACACCTTCTACAATG | This study. |
| *ACTB RP* | AATGTCACGCACGATTTCCCGC | This study. |
| *FOXP1 FP* | CAGATATTGCGCAGAACCAA | https://molecularautism.biomedcentral.com/articles/10.1186/2040-2392-4-23 |
| FOXP1 *RP* | GCAAACATTCGTGTGAACCA | https://molecularautism.biomedcentral.com/articles/10.1186/2040-2392-4-23 |
| ***Others*** | | |
| mTeSR1 | STEMCELL Technologies | Cat#252050 |
| Gentle Cell Dissociation Reagent (GCDR) | STEMCELL Technologies | Cat#100-0485 |
| Accutase | Sigma aldrich | Cat#A6964-100ML |
| DMEM/F12 | ThermoFisher | Cat#11330032 |
| KnockOut Serum Replacement (KSR) | ThermoFisher | Cat#10828-028 |
| Penicillin-streptomycin Solution | ThermoFisher | Cat#15140122 |
| GlutaMAX | ThermoFisher | Cat#35050061 |
| NEAA (MEM Non-Essential Amino Acids Solution) | ThermoFisher | Cat#11140050 |
| β-mercaptoethanol | ThermoFisher | Cat#21985023 |
| Heparin | STEMCELL Technologies | Cat#7980 |
| Ultra-low attachment U-bottom 96-well plates | Sbio | Cat#MS-9096UZ |
| Ultra-low attachment 24-well plates | Sbio | Cat#MS-90240 |
| Ultra-low attachment 6-well plates | Sigma aldrich | Cat#CLS3471-24EA |
| Neurobasal Medium | ThermoFisher | Cat#21103049 |
| B-27 (minus vitamin A) | ThermoFisher | Cat#12587010 |

**Supplementary Table 2. Summary of therapeutic modalities on nGD MLOs.**

| | | GCase activity (% of WT, Mean ± SEM) | | | Substrate GluSph level (Fold to WT, Mean ± SEM) | | | TH expression (Fold to WT, Mean ± SEM) | | |
|---|---|---|---|---|---|---|---|---|---|---|
| **iPSC line derived MLO** | **Therapeutic modalities** | **w/o treatment** | **w/ treatment** | **P value** | **w/o treatment** | **w/ treatment** | **P value** | **w/o treatment** | **w/ treatment** | **P value** |
| **GD2-1260** | CRISPR/Cas9-mediated gene correction | 15.9% ± 2.0% Wk8 | 46.3% ± 1.8% Wk8 | P<0.001 | 6.6 ± 1.6 Wk28 | 1.4 ± 0.3 Wk28 | P<0.05 | 0.24 ± 0.02 Wk8 | 1.06 ± 0.19 Wk8 | P<0.001 |
| | SapC-DOPS-fGCase (2 weeks treatment) | 5.6% ± 0.5% ~Wk15 | 95.2% ± 2.1% ~Wk15 | P<0.001 | 3.9 ± 0.8 Wk16 | 0.7 ± 0.1 Wk16 | P<0.001 | N/A | N/A | N/A |
| | AAV9-GBA1 (3 weeks treatment) | 6.0% ± 0.6% Wk16 | 47.8% ± 2.3% Wk16 | P<0.001 | 3.1± 0.4 Wk16 | 0.3 ± 0.03 Wk16 | P<0.001 | 0.65 | 0.73 | n.s. |
| | GZ452 | N/A | N/A | N/A | 5.9 ± 0.8 Wk15 | 1.1 ± 0.03 Wk15 | P<0.001 | N/A | N/A | N/A |
| **GD2-10-257** | CRISPR/Cas9-mediated gene correction | 8.8% ± 0.2% ~Wk15 | N/A | N/A | 3.6 ± 0.02 Wk16 | N/A | N/A | 0.20/Wk16 0.17/Wk28 | N/A | N/A |
| | SapC-DOPS-fGCase (2 weeks treatment) | 8.8% ± 0.2% ~Wk15 | 88.0% ± 9.1% ~Wk15 | P<0.001 | 7.6± 0.9 Wk16 | 3.3 ± 1.0 Wk16 | P<0.001 | N/A | N/A | N/A |
| | AAV9-GBA1 (3 weeks treatment) | 8.8% ± 0.2% Wk16 | 37.7% ± 4.0% Wk16 | P<0.001 | 8.4 ± 0.5 Wk16 | 0.6 ± 0.1 Wk16 | P<0.001 | N/A | N/A | N/A |
| | GZ452 | N/A | N/A | N/A | N/A | N/A | N/A | N/A | N/A | N/A |

N/A, not available; n.s., not significant. GluSph, glucosylsphingosine.

**Supplementary Table 3. Dysregulated pathways in nGD models.**

| Model | Tissue | Key Dysregulated Pathways |
|---|---|---|
| Human nGD organoid | Midbrain-like organoids | Nervous system development<br>Axon guidance<br>Neuron differentiation<br>Dopaminergic/Glutamatergic/GABAergic synapse<br>Apoptosis etc. |
| Mouse nGD model[#] | Brain (Midbrain region) | Neurological disease<br>Axonal guidance signaling<br>Dopamine/Glutamate/GABA receptor signaling<br>Lipid metabolism<br>Cell death and survival etc. |

#, Data from "Dasgupta, N., Xu, Y.H., Li, R., Peng, Y., Pandey, M.K., Tinch, S.L., Liou, B., Inskeep, V., Zhang, W., Setchell, K.D., Keddache, M., et al. (2015). Neuronopathic Gaucher disease: dysregulated mRNAs and miRNAs in brain pathogenesis and effects of pharmacologic chaperone treatment in a mouse model. Human molecular genetics 24, 7031–7048. 10.1093/hmg/ddv404." (https://pubmed.ncbi.nlm.nih.gov/20047948)
